# Stress-mediated growth determines *E. coli* division site morphogenesis

**DOI:** 10.1101/2024.09.11.612282

**Authors:** Petr Pelech, Paula P. Navarro, Andrea Vettiger, Luke H. Chao, Christoph Allolio

## Abstract

In order to proliferate, bacteria must remodel their cell wall at the division site. The division process is driven by the enzymatic activity of peptidoglycan (PG) synthases and hydrolases around the constricting Z-ring. PG remodelling is regulated by de-and re-crosslinking enzymes, and the directing constrictive force of the Z-ring. We introduce a model that is able to reproduce correctly the shape of the division site during the constriction and septation phase of *E. coli*. The model represents mechanochemical coupling within the mathematical framework of morphoelasticity. It contains only two parameters, associated with volumetric growth and PG remodelling, that are coupled to the mechanical stress in the bacterial wall. Different morphologies, corresponding either to mutant or wild type cells were recovered as a function of the remodeling parameter. In addition, a plausible range for the cell stiffness and turgor pressure was determined by comparing numerical simulations with bacterial cell lysis data.

## Introduction

Bacteria are single-cell organisms that multiply by binary fission. Unlike most eukaryotic cells, bacteria are enveloped by multiple layers.[1] The hardest of these layers, protecting the cell from osmotic lysis, is the bacterial wall, which is formed of peptidoglycan (PG), a peptide-crosslinked glycan polymer. Hence, unlike cell membranes, the bacterial wall cannot be divided by purely mechanical forces generated by protein assemblies. Instead, bacterial division requires the enzymatic activity of a combination of precisely balanced PG synthases and hydrolases. Inhibition of this process is a major target of antibiotics[2–5], as they are specific to bacteria. In recent years our understanding of bacterial growth and division has been enhanced greatly[6–12]: Bacterial genetics studies have identified the determining factors for cell wall synthesis, homeostasis and division. While the detailed mechanisms of how the different proteins in the divisome mediate cell division remain not fully understood, cryo-electron tomography (cryo-ET) was able to furnish the ultrastructure of the division site of *E. coli* as a function of PG synthesis and hydrolysis levels, thereby providing a basis for quantitative analysis of its division mechanism.[13]

As of yet, there has not been any successful model of the division mechanism that faithfully reproduces division site ultrastructure and morphogenesis. Moreover, for such a model to be useful for interpreting and predicting the behavior of the cell, it should map genetic modifications to observed morphology via a transparent mechanism. We are not aware of any published attempts to meet the latter challenge. The model should also be physical and parsimonious, which excludes purely phenomenological approaches. To wit, older models often resulted in reasonable morphologies of the bacterial surface, but the constriction dynamics evolved independently of any cell process; moreover, PG was described as thin shell and so, structural details, such as septal wedge formation could not be modeled.[14, 15] Recent coarse-grained particle-based models[16–18] have introduced a stylized mechanistic understanding, but also lack realistic material properties and are unable to describe the formation and structure of bacterial septa. *E. coli*growth and division are responsive to mechanical stimuli in a way that suggests mechano-chemical coupling of the growth apparatus.[19] Models of stress-mediated growth are often formulated within the mathematical framework of morphoelasticity[20]. They have been widely used to describe growth in macroscopic systems.[21–27] Morphoelasticity combines large-strain elasticity with the possibility of a permanent change of volume and shape. This change is attributed to growth and remodeling, respectively. Morphoelasticity is closely related to the continuum description of elastoplastic deformations, with the main difference being that the plastic deformation can change the volume, hence it becomes a description of growth. Gradual growth leads to residual stresses in the body, which in turn influences material restructuring and deposition. Constructing a morphoelastic model of bacterial division of *E. coli* has the potential to greatly enhance our understanding of the mechanobiology of the bacterial cell envelope. In contrast to stress-mediated growth in a multicellular organism, it is even possible to employ simple thermodynamic considerations to explain how stress is related to growth at the molecular scale. The resulting model, yields a microscopic and quantitative description of the forces acting in each stage of cell division and the way they determine the ultrastructure of the division site. Our description is also the first to take into account the varying thickness of the material.

## Results and Discussion

### Turgor Pressure and Mechanical Properties of Peptidoglycan

Accurate knowledge of elastic properties and traction forces is crucial for any model, in which growth is mediated by stress. Unfortunately, there is neither a consensus on the mechanical properties of the bacterial wall nor on the size of the turgor pressure. The bacterial wall consists of PG, which is made of up to *∼* 200 nm long glycan strands[28], whose average length is *∼* 35 nm[29]. These glycan strands are crosslinked by short peptides (Fig. 1a). In *E. Coli*, most of the bacterial wall consists of a single PG layer with a thickness of between *∼* 2.5 nm and *∼* 6 nm.[28–32]

**Fig. 1.**
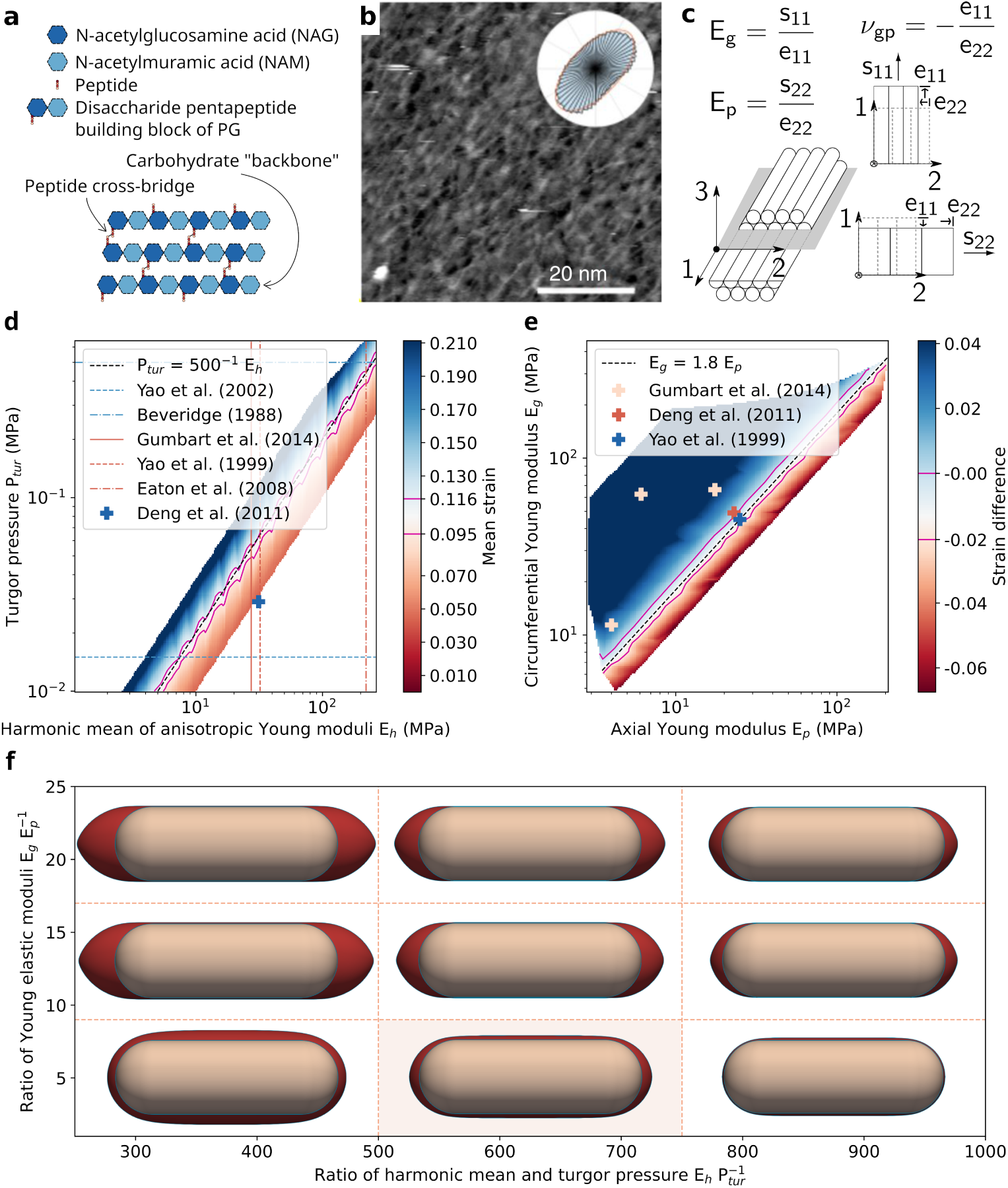
PG structure and elastic properties. **a**, Schematic of PG crosslinked structure. **b**, Prevalent fiber orientation. From [28] under CC 4.0 (http://creativecommons.org/licenses/by/4.0/) **c**, Three elastic moduli, associated deformations of a transversely isotropic bulk material. **d**, Mean strain *εm* observed in numerical simulations shown in a log-log plot of the turgor pressure *Ptur* and the cell’s stiffness *E_h_*. A 10% interval around the experimental value (*ε_m_ ≈* 0.1) is indicated by blue lines. Linear relation traced by dashed black line. Experimentally measured values of either *E_h_*or *P_tur_* marked by lines, simultaneously measured values by crosses. **e**, Strain difference *ε_d_* observed in numerical simulations, shown in a log-log plot of axial and circumferential Young’s modulus *Ep* and *E_g_*. A 100% interval around the experimental value (*ε_d_ ≈* 0.01) is marked by blue lines. Correlation depicted by dashed black line. Experimentally measured values marked by crosses. **f**, Numerically computed shapes of pressurised sacculi (maroon) compared to their lysed state (beige). Shape with experimentally observed *ε_m_*and *ε_d_* displayed on a light beige background.

Although the PG structure is disordered, and strands partially overlap[28], a preferential orientation of the glycan chains along the circumferential direction of the bacterial rod is widely assumed. In contrast to this, the fiber orientation at the poles is not clearly established.[28] To illustrate the situation, we show the anisotropy data as measured away from the poles by Turner et al.[28] in Fig. 1b. Consequently, the material properties of PG are expected to be anisotropic, and have been reported as such.[32]

The partially disordered long-range structure of PG complicates any attempt to estimate its elastic parameters in atomistic simulations.[33] Direct experiments on living cells are also difficult to interpret, as a living organism contains several active and passive regulatory mechanisms.[34] Therefore, estimates of the Young’s modulus of PG differ by several orders of magnitude, from 5 *·* 10*^−^*^2^ to 2 *·* 10^2^ MPa, even with state-of-the-art techniques, such as atomic force microscopy (AFM).[35–37] While examining sacculi of dead cells increases control over the experimental conditions, differences in sample preparation might strongly affect results as PG is known to be very sensitive, e.g. to hydration.[32]

To determine the magnitude of the turgor pressure, several techniques have been used: collapse of gas vesicles (in other Gram-negative bacteria) [38, 39], AFM indentation, [40, 41], and estimation of the total chemical content of the cytoplasm [42]. The estimated pressure values vary by more than an order of magnitude, from 0.1 to 3 bar. AFM indentation is a direct probe, yet it is difficult to separate the internal pressure from the elastic wall contributions; values obtained in such a way hence represent only an upper bound on the real turgor pressure. A lower bound is provided by a study of a membrane bulge on mutated *E. coli* [43], where the PG was intentionally weakened and damaged to allow formation of a bulge, and consequently the bacteria may have reduced their internal pressure via some safety mechanism.

Given the large uncertainty about the range of the turgor pressure and elastic moduli of PG in particular, we decided to use an independent experimental measurement for cross checking plausible values: We estimated the strain in a living cell relative to its unpressurized state using a plasmolysis experiment as a reference.[44] In order to test plausible values for elastic parameters, we computed the deformation of the cell wall from a stress-free reference configuration, as shown in green in Fig. 1f. Then, we compared its strain with the experimental values for a given set of elastic moduli and turgor pressure. In accordance with the plasmolysis experiment we set the length and width of an unpressurized sacculus to 3 *µ*m and 1 *µ*m respectively. Our reference strains for the turgor pressure are those reported for *E.Coli* in lysogeny broth, namely the longitudinal strain *ε_l_ ≈* 12 % and the radial strain of *ε_w_ ≈* 10 % [44]. From the range of experimentally observed values, we chose the thickness of 6 nm[32, 43].

To compute the deformation, we used an anisotropic, nonlinear elastic model and solved it with the finite element method (see the SI for more details on the model and its numerical implementation). In agreement with fiber orientation, the material response differs in the circumferential and axial direction. This corresponds to a transversely isotropic bulk material, see Fig. 1c for a schematic of the material structure and response. Such a material has five elastic moduli, however, as the PG is treated as a thin layer, only three are considered in experiments and simulations[32, 33]; the Young’s modulus and Poisson ratio in the direction normal to the surface are neglected. The three remaining parameters are two Young’s moduli, the axial *E_p_* and circumferential *E_g_*, and one Poisson ratio *ν_gp_* = 0.48, which we took from the only available measurement by Yao et al.[32]. This value of *ν_gp_*is also comparable to MD calculations by Roux et al.[33]. Overall, the parameters indicate that the thin PG layer has low compressibility.

Summarizing, the elastic parameters that need to be consistent with the lysis experiment are the two Young’s moduli *E_g_* and *E_p_*. Their harmonic mean *E_h_* = 2*E_g_E_p_*(*E_g_* +*E_p_*)*^−^*^1^ defines the material’s overall stiffness and their ratio *E_g_E_p_^−^*^1^, by dif-fering from 1, expresses its anisotropy. In order to facilitate the comparison we defined corresponding strain measures; from the longitudinal strain *ε_l_*and the radial strain *ε_w_* we computed the mean strain *ε_m_* := ½(*ε_l_* + *ε_w_*), describing the total strain of the bacterial wall, and the strain difference *ε_d_* := *ε_l_ − ε_w_*, which measures the anisotropy of the deformation. In Fig. 1d we show simulation values of *ε_m_* in a log-log plot of the stiffness *E_h_* against the turgor pressure *P_tur_*, and in Fig. 1e we show numerically computed values of *ε_d_* in a log-log plot of the Young’s moduli *E_h_* and *E_p_*, both containing values reported in the literature. An early review by Beveridge (1988)[45] estimates the turgor pressure in the range 3 *−* 5 atm, referring e.g. to nephelometric experiments on bacteria with gas vesicles[38]. Later atomic force microscopy (AFM) measurements by Yao et al. (2002)[41] suggest values 0.1 *−* 0.2 atm (for the related Gram-negative bacterium *Pseudomonas aeruginosa*). Elastic moduli were measured via AFM by Yao et al. (1999)[32] on sacculi of dead cells and by Eaton et al. (2008)[37] on live bacteria. MD simulations of a PG matrix segment were performed by Gumbart et al. (2014)[33]. An AFM experiment determining separately turgor pressure and elastic moduli was performed by Deng et al. (2011)[43].

The comparison shown in Fig 1d and e reveals that when both the strain measures *ε_m_* and *ε_d_*lie close to their experimental values, the stiffness *E_h_* correlates with the turgor pressure *P_tur_* while the Young’s modulus *E_g_*correlates with *E_p_*. If the value of the turgor pressure *P_tur_* is known, the cell wall’s stiffness is determined from the plot by *E_h_ ≈* 500 *P_tur_* (see the SI for further details) and the Young’s moduli by *E_g_ ≈* 1.8 *E_p_*. Moreover, since both the values of the strains *ε_l_* a *ε_w_* lie around 0.1, the material response is close to a linear regime.

It has been reported that the PG matrix stiffens as its strain increases, i.e. the elastic moduli estimated via Hertzian contact mechanics on a living, pressurized bacterium scale (almost linearly) with the turgor pressure[43, 46]. As the turgor pressure is believed to be 10 times higher for cells treated in a distilled water than for cells in a growth medium[41], this scaling may help to explain the discrepancy between studies on the PG’s elasticity[32, 36, 37, 43]. Since our model departs from a stress-free state (i.e. unpressurized sacculus to which the turgor pressure is then applied), we take the values from [32], the only literature source providing elastic moduli of stress-free sacculus. These elastic moduli are then compatible with a moderate turgor pressure.

To summarize, we consider in our model a sacculus of length 3 *µ*m, diameter 1 *µ*m, and thickness 6 nm in its unpressurized state; as discussed above, these values agree with experiments, are internally consistent and have been used in other studies[32, 43]. We set *P_tur_* = 0.6 atm = 6 *·* 10*^−^*^2^ MPa as then the PG stiffness *E_h_* determined from the plot Fig. 1d agrees with the values from [32]. This value of the turgor pressure provides consistency with the more recent elastic measurements and the lysis data. The compatible elastic parameteres are hence *E_g_ ≈* 42 MPa and *E_p_ ≈* 23 MPa. As already mentioned, we consider the Poisson ratio *ν_gp_* = 0.48.[32, 33]

As PG is stiffer than the outer and inner membrane, we do not include lipid membranes into the model and when referring to them either the exterior or interior of the sacculus is meant. We exploit the known (approximate) cylindrical symmetry of the rod-like shape of *E. coli* (which is maintained in the division process) in our model.

### Cell Division by Stress-driven PG Growth and Remodeling

The multiprotein assembly called divisome directs PG synthases and hydrolases, which build and remodel the PG matrix at the division site. Its key components are 1. the *Z-ring* (whose formation is directed by the tubulin-like FtsZ protein) 2. the *PG synthesis* complexes (FtsW and FtsI synthase[47, 48] regulated by the FtsQLB and FtsN complexes)[49], and 3. the *PG hydrolysis* by amidase enzymes (activated by the FtsEX-EnvC complex)[50, 51].

PG precursors are synthesized in the cytoplasm and then flipped across the inner membrane to face the periplasm where they are subsequently incorporated into the PG matrix by the PG synthase FtsWI. A portion of already existing PG is modified by amidases that remove the peptide crosslinks from the glycan chains. The rate of septal PG (volume) production estimated by fluorescence measurements is of the order 10^3^ nm^3^ *·* s*^−^*^1^.[13] In our model, the cell has at the beginning a length *L* = 3 *µ*m, width *W* = 1 *µ*m, and thickness H := R_out_ *−* R_in_ = 6 nm (see Fig. 2a);

**Fig. 2.**
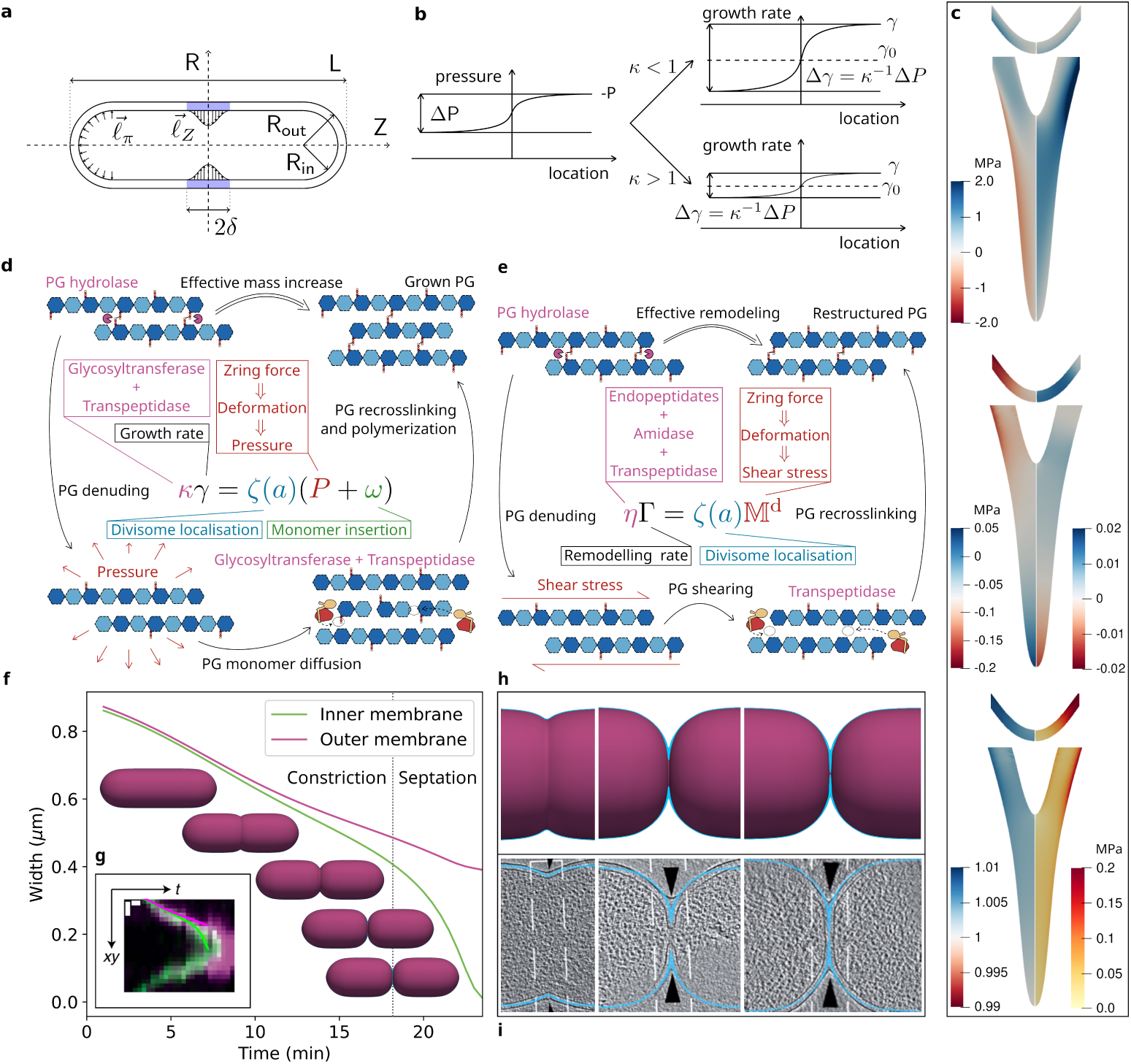
Stress-mediated remodeling and growth during the division of a wild type *E. coli* and its comparison with experiment[13]. **a**, Layout of the sacculus with depicted parameters and pressures. **b**, Influence of the pressure profile on the resulting growth rate. **c**, Panels from top to bottom: First: Detail of the growth zone with a distribution of the shear stress (left) and the normal radial stress (right) during the constriction phase (top) and the septation phase (bottom). Second: Detail of the active zone with a distribution of the pressure (left) and the growth rate (right) during the constriction phase (top) and the septation phase (bottom). Growth rate is a dimensionless quantity. Third: Detail of the active zone with a distribution of the relative density (left) and energy density (right) during the constriction phase (top) and the septation phase (bottom). Relative density, as a ratio of the current over the one at relaxed, unstressed state, is dimensionless. **d**, Schematic of the effective remodeling. **e**, Schematic of the effective growth. **f**, Radii of the inner membrane (IM) in green and the outer membrane (OM) in purple at the division site. Several stages of the cell division shown below the curves. Transition between the constriction and septation phase is set to the moment when IM narrows two times faster than OM. **g**, Curves from the plot in (**f** ) overlaid with experimental kymographs, measured via fluorescently labeled membranes at the division site[13], with depicted radial (xy) and temporal (t) directions. **h**, Detailed view of the division site predicted by the model at selected times; OM in purple, PG cut highlighted in blue. **i** PG cut in blue as predicted by the model overlaid with cryo-electron tomography data[13].

### Z-Ring Force

The Z-ring is composed of tubulin-like FtsZ filaments (app. 100*−*200 nm in length[52]) which recruit the PG synthesizing/cleaving apparatus to constrict the cell. Each FtsZ monomer is about 4 nm long. Upon GTP hydrolysis, it bends and contributes 20 *−* 30 pN to the constriction force, as estimated from MD simulation.[17] The number of monomers is believed to lie between 5000 and 7000 in the whole *E. coli* cell from which 30 *−* 40% is incorporated into the Z-ring and the rest is cytoplasmatic[53, 54]. The *∼* 2100 FtsZ molecules in the Z-ring could form a total protofilament 8400 nm long, which would encircle a 1-*µ*m-diameter cell two and a half times. Hence, we expect the Z-ring constriction force line density in units to a few tens of pN *·* nm*^−^*^1^. The Z-ring width stays approximately at 84 *±* 2 nm during the whole division.[52]

The turgor pressure *ℓ_π_* expands the cell everywhere from inside while the Z-ring pressure *ℓ_Z_* constricts it within the active zone, being highest in the middle and vanishing at the distance *δ* = 45 nm. The Z-ring force line density *f_Z_* points inwards along the circumference and is given as a line integral of the Z-ring pressure *ℓ_Z_* along the axial cross-section of the inner membrane within the *δ* distance (Fig. 2a), i.e. 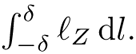. In other words, the Z-ring force applies up to an arc-length of *δ* along the inner boundary of the bacterial wall. Modeling the Z-ring as a mechanical force[15, 16], locating it via the arc-length of the shell boundary[15] are natural steps. Following the literature data mentioned above we set *f_Z_* = 6 pN *·* nm*^−^*^1^ and *δ* = 45 nm.

### Stress-mediated Growth

The orientation of the PG glycan strands was found to be far from perfectly uniform and periodic, see the AFM data with the distribution of fiber orientations in Fig. 1b (taken from [28]). Although it is in principle possible to account for this in detailed simulations (e.g. by a distribution of orientations or the degree of crosslinking), doing so would increase the number of unknown parameters of the problem. Given the lack of data on these quantities available from the literature, we decided rather to reduce the number of necessary parameters to an absolute minimum. Hence, we use a morphoelastic model of PG growth and remodeling within continuum mechanics[55, 56] with constant mechanical properties, including an effective circumferential orientation of fibers. The phenomenology of remodeling and growth emerges from intricate feedback loops of enzyme kinetics, protein activity, and diffusion happening at the molecular scale. To build a useful physical model of the mechanochemical process we need to remove degrees of freedom.[57] The elastic relaxation time of the cell wall is *∼* 1 s[58] and the division time is of the order of *∼* 10^3^ s in the laboratory under favorable conditions[59]. This separation of timescales allows us to employ a quasistatic approximation to modeling PG growth, i.e. neither inertial nor viscous effects are important. In this context, we can assume these rates to depend mainly on the chemical potentials of proteins and substrates, and in particular deduce a similar dependence of all chemical potentials on the stress (see the SI).

Overall, our model is a variant of what is known in the literature as stress-mediated growth which is empirically justified for many systems: muscles, bones or trees[22–25] For example, it is well established that turgor pressure in plant cells drives mechanical expansion of the cell wall during growth[60, 61]. Also *E. coli* cells, when exposed to constant bending forces for an extended period of time, grow into curved shape which they maintain even after unloading[19].

### PG Growth and Insertion

The growth of PG via precursor insertion leads to local volume increase, local volume decrease is associated with PG recycling and hydrolysis. The global rate of septal PG volume increase Ω was measured to be *≈* 10^3^ nm^3^ *·* s*^−^*^1^.[13] As the cell wall thickness is on the order of nm, the area increase is ca. *∼* 10^3^ nm^2^ *·* s*^−^*^1^. This increase is much slower than the diffusion of PG precursors as the diffusion coefficient of periplasmic proteins is *∼* 3 *·* 10^6^ nm^2^ *·* s*^−^*^1^[62] The diffusion in the porous environment allows the precursors to attain equilibrium distribution, consistent with our quasistatic approach. The porosity of the PG medium arises as e.g. PG is processed by cell wall hydrolases. Hydrostatic pressure can then pull some of the strands apart, enlarging the gap between molecules. This in turn eases the entry of PG precursors into the area. Growth is then accomplished by PG synthases, prominently FtsWI, integrating the new material into the matrix (see Fig. 2e). It follows that the growth rate *γ* should depend on the pressure. In our description, the pressure *p* has to be scaled to account for the elastic deformation. The rescaled pressure field is called *P* . As the FtsWI are known to be collocated with the divisome, we introduce a localizing function *ζ* (similar to a Gaussian distribution) to restrict the growth to the environment of the Z-ring. (for further details see the SI). The total net volume of PG units incorporated into the matrix per time (denoted Ω) is treated as a fixed parameter of the model and hence the mean value of the growth rate is fixed via the Lagrange multiplier *ω*. As it is easiest to assume a linear relation (and in the end completely sufficient), we introduce a proportionality constant for the growth rate, the resistance modulus *κ*. As the mean value is already fixed, it determines only the sensitivity of the growth rate *γ* to the pressure difference Δ*p*. Low resistance *κ <* 1 leads to a growth rate that varies more over the active zone, while high resistance *κ >* 1 makes the growth rate almost constant across space (see Fig. 2b). We associate the sensitivity with the presence and concentration of glycosyltransferase and transpeptidase enzymes. The change of PG volume is not only due to the incorporation of new material, but also due to the recycling and digestion of existing PG. Hence, the concentration of mobilized material is also a function of the remodeling apparatus. For example, if a lot of material is available for incorporation, the insertion depend only on the localization of the divisome. At the same time, the Lagrange multiplier *ω* shifts the curve in such a way that the total volume rate matches the prescribed value. Finally, when the arc-length distance *a* from the middle of the bacterium is larger then the active zone radius *δ*, the localization function *ζ* vanishes and no growth occurs. The whole growth law is given in Fig. 2e, together with a color-coded schematic intended to illustrate the underlying processes.

We consider the net volume rate of septal PG incorporation Ω = 10^3^ nm^3^ *·* s*^−^*^1^[13]. Numerical experiments show that *κ* has almost no influence on the division process; we set *κ* = 44 MPa . s.

### PG Remodeling

The degree of crosslinking and re-crosslinking, given by enzyme activity, allows for sliding of the polymer strands over each other. This relaxation does not result in a volumetric change and is driven by the stress inside the material. Namely, by a ”pressure-free” stress, the so-called deviatoric part of the Cauchy stress. As for the pressure, also the Cauchy stress field has to be transformed, such that it accounts for the elastic deformation. We denote the resulting quantity M*^d^*^1^ .

The solid PG structure is locally disrupted by hydrolysis of some of the peptide crosslinks. The shear stress carried by them is then redistributed to the rest of the matrix as the denuded PG strands slide over each other until a new equilibrium position is reached. The PG chains are finally re-crosslinked by transpeptidases, reestablishing a stiff structure. The more enzymes are active, the more denuding occurs and the more the structure will comply to the shear stress, which in turn leads to a faster remodeling. The PG effectively behaves as a melted polymer composed of reptating chains. As it is common in polymer science[63–66] the remodeling rate matrix Γ (3 *×* 3) is proportional to the ‘driving force’ M*^d^* and we call the corresponding modulus *η* ‘solidity’. When the arc-length distance *a* from the middle of the bacterium is larger then the active zone radius *δ*, the localization function *ζ* vanishes and no remodeling occurs. The equation for Γ and the relevant biochemical processes are illustrated in Fig. 2d.

The solidity *η* is the crucial parameter in our model that reflects enzyme activity and specifies the balance between growth and remodeling, which in turn influences the cell’s morphology at the division site.

### Phenomenology of the Division Process

At the beginning of the division process, the inward pulling Z-ring force induces two significant stresses: shear stress deflecting the sacculus to the center of the cell, resulting in a bending deformation, and normal stress in the radial direction expanding its thickness. In this discussion thickness is measured by the extension of PG in the radial direction. The distribution of these stresses is shown in Fig. 2c. Predictions of our model are determined by the fixed mass increase of PG. The location of mass increase is then stress-mediated, i.e. the growth and the remodeling of the material try to reduce stress by directing material modification to follow it. Consequently, the PG growth and remodelling of the cell wall gradually bends it inwards and thickens it from the inside. During the constriction, it eventually forms a small wedge of new sPG. As the division site changes its shape, the shear stress decreases while the normal stress in the radial direction increases. The thicker the PG at the division site gets, the more the normal stress concentrates in the middle of the wedge, as the arc-length restriction will limit the Z-ring and the growth zone onto the forming septum. Hence, instead of bending the whole structure as in the beginning, the Z-ring force rather strains the wedge, which directs the elongation via the PG synthases into the radial direction. This positive feedback loop leads in the end to a rapid closure of the septum.

The growth rate distribution (shown on the right side of the middle panel of Fig. 2c) is in the initial stage homogeneous across the thickness, highest in the middle of the cell and decreases smoothly with the arc-length distance from the center of the Z-ring. In this way it models the collocation of the growth and remodeling apparatus with FtsZ. Later, approximately when the cell wall in the midplane is twice as thick as at the beginning, the domain of maximum growth rate leaves the precise center and moves further aside; also the growth rate is higher near the inner membrane and lower at the outer PG boundary. This distribution remains roughly unchanged till the end of the septation phase, the highest mass increase thus occurs at the ‘neck’ of the septum. Also, as the volume in which the growth happens increases, due to the constraint the growth rate spatial density spreads out and its peak decreases.

Roughly speaking, the PG in the active zone behaves as an elastic, almost solid glue which, as it is being pulled inside, slowly forms a small drop; however, as new PG is continuously incorporated, no necking occurs, the drop does not separate from the cell wall and instead forms the septum. This process also can lead to the wedge-shapes observed in experiment. The septum hence grows rather at the neck from where is being pulled down by the Z-ring. In particular, the septum does *not* grow from its tip. While this finding is in contrast with what is widely assumed in the field, it should be pointed out that the width of the growth zone itself is equal to that of the Z-ring, and is 90 nm, a value close to what is given in the literature.[15, 16] This implies that the Z-ring is wrapped around parts of the septum. Our model reflects the dimensions and position uncertainty of the growth apparatus.

### Quantitative Model of Wild Type Cell Division

The correct morphology of the division site during the division (i.e. the constriction and septation phase, which we model) of the wild type bacteria was recovered for *η* = 66 MPa *·* s, *κ* = 44 MPa *·* s, and Ω = 10^3^ nm^3^ *·* s*^−^*^1^. The predictions of the model and their comparison to published cryo-ET and fluorescence microscopy data[13] are shown in Fig. 2f-i. We observe an initially slow constriction, which is followed by a short and accelerating septation (Fig. 2f). This observation is in quantitative agreement with experimental kymographs, obtained by fluorescently labeling inner and outer bacterial membrane (Fig. 2g). Also the final architecture of the division site and the width and thickness of the septum are faithfully recovered, as can be seen by overlay with electron micrographs (Fig. 2h,i); the full original micrographs are included in SI, see the Methods section for further details. The (unrescaled) pressure field *p* is depicted in the middle panel of Fig. 2c, relative density (the ratio of the current density over the referential at the relaxed, unpressurized state) together with the energy density are shown in the bottom panel of Fig. 2c.

### Modelling of Mutant Strains

The Δ*envC* -mutant is defective for one of two amidase activators[67]. Reduced amidase activity corresponds in our model to higher solidity *η*, because PG remodelling is suppressed. We hence decided to check the generality of our model by trying to capture the division of mutated cells by altering the solidity *η* and resistance *κ*. The results for *η* = 93 MPa *·* s, *κ* = 62 MPa *·* s, and Ω = 10^3^ nm^3^ *·* s*^−^*^1^ are shown in Fig. 3a-d. As for the wild type cells, the septation starts very progressively at the late stage of the division both in the model and experiment (Fig. 3a-b). However, for this mutant, the constriction deformation is severely impaired, so that the septation is almost ”vertical”. This follows naturally from the suppression of PG remodelling, which is necessary to achieve bending-like behavior. The resulting large contact surfaces of the flat septa experimentally lead to a failure of separation of the bacteria, and to the formation of filament-like colonies[13]. The model also predicts properly the two main differences in the mutant’s division. Namely, the wider septum with a proper shape (Fig. 3c-d) and the longer division time which is in average two times longer than for the wild type (Fig. 3g), also reported earlier[13].

**Fig. 3.**
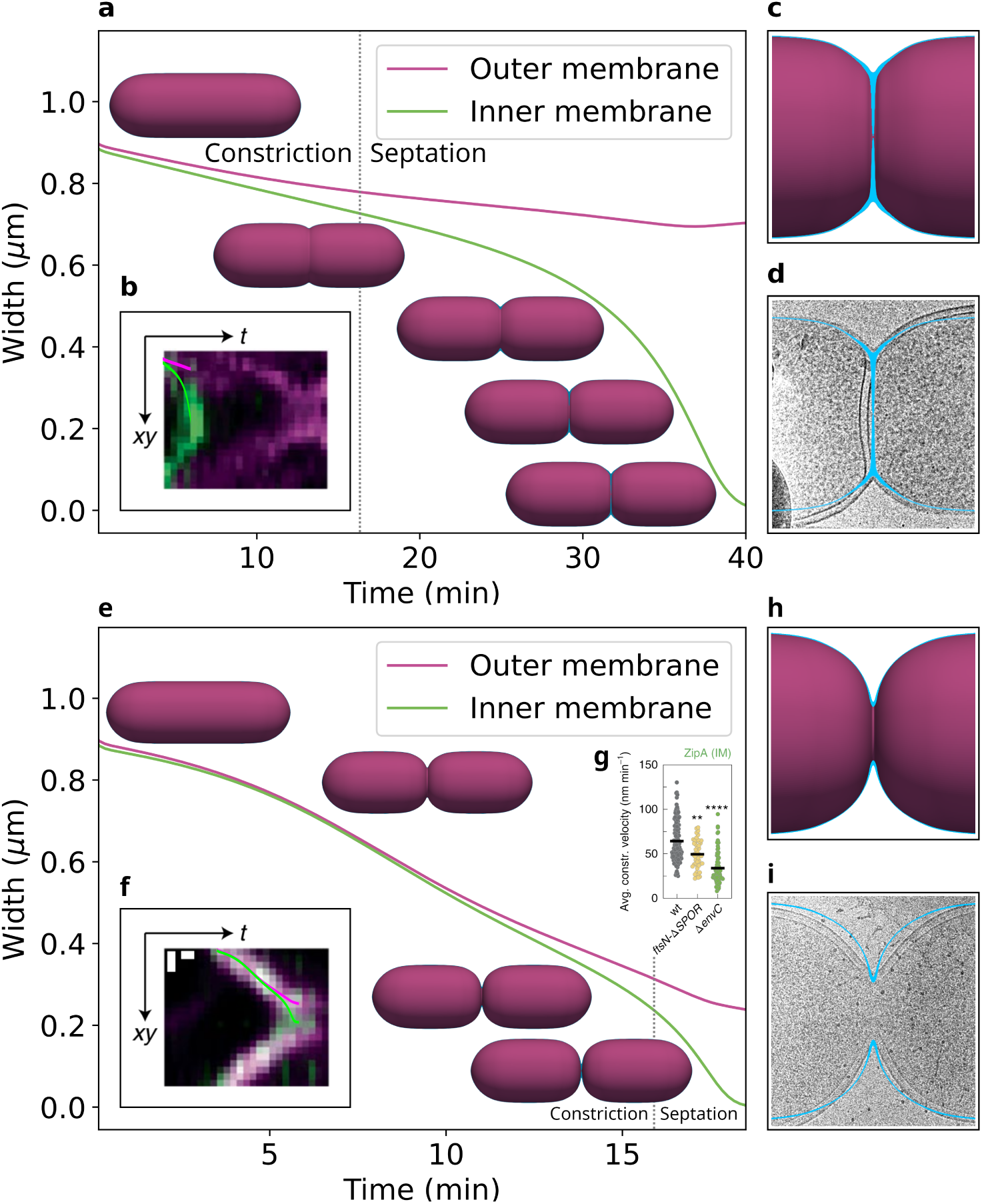
Stress mediated remodeling and growth during the division of the Δ*envC* -(top) and *ftsN-*Δ*SPOR*-(bottom) mutants of *E. coli* and their comparison with experiment. **a**, **e**, Radii of the inner membrane (IM) in green and the outer membrane (OM) in purple at the division site. Several stages of the cell division shown along the curves. **b**, **f**, Curves from the plot in (**a**) and (**e**) respectively overlaid with experimentally measured positions of the membranes at the division site[13], with depicted radial (xy) and temporal (t) directions. **c**, **h**, Detailed view of the division site predicted by the model at a selected time; OM in purple, PG cut highlighted in blue. **d**, **i** PG cut in blue as predicted by the model overlaid with cryo-electron tomography images. **g**, Average constriction velocities measured in experiment[13]. Brown–Forsythe and Welch ANOVA test with Dunnett’s correction for multiple comparisons, significance of differences is tested relative to wild type (wt); *^∗∗^P <* 0.01, *^∗∗∗∗^P <* 0.0001; *N* = 150 (wt), 48 (*ftsN-*Δ*SPOR*), 74 (Δ*envC* ) kymographs.

The *ftsN-*Δ*SPOR*-mutant lacks the SPOR domain that recognizes the denuded PG and thereby activates the FtsQLB complex. As this suggests reduced incorporation of PG precursors, we tried to model this mutant with reduced volume rate Ω = 900 nm^3^ *·*s*^−^*^1^, while keeping the solidity and resistance moduli same as for the wild type *η* = 66 MPa *·* s and *κ* = 44 MPa *·* s. The results are shown in Fig. 3e-i. Unlike the wild type cell, the OM of the *ftsN-*Δ*SPOR*-mutant constricts for most of the division at a similar rate as the IM and the much narrower septum is formed quickly at the latest stage of the process; this behaviour is in an agreement with experimental data[13], see Fig. 3e,f,h,i. Although the constriction and septation agree with experimental data and the final shape is faithfully recovered, mere reduction of the volume rate counting for reduced PG insertion is not enough, as the *ftsN-*Δ*SPOR*-mutant should divide slightly slower than the wild type cell (see Fig. 3g). Nevertheless,we show in the next section, the three parameters *η*, *κ*, and Ω can be scaled properly such that the division runs along the same lines, while the total division time can be altered.

Full original micrographs of both mutants are included in SI, see the Methods section for further details on image processing. Note, that Cryo-EM images are necessarily of different individual cells and that these cells do not have perfect symmetry due to technical limitations inherent to the method. Considering these limitations, our model agrees with the data.

### Sensitivity of the Model to Parameter Variations

The model’s predictions are stable with respect to the geometric parameters. Varying the length or width of the bacterium in the order of a few tenths of its value leads only to a minor quantitative change of the PG matrix’ elasticity, remodeling, and growth. When reducing the cell wall’s thickness, the elastic moduli have to be increased to maintain the bacterial shape during division, because the elastic response is governed by a product of these. In order to initiate the constriction phase it is necessary that the Z-ring force should be strong enough to overcome the turgor pressure, i.e. approximately

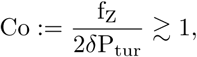

where 2*δ* is the width of the Z-ring. We call Co the constriction number. For Co *<* 1 the cell in the middle expands in radius and its wall bulges, while Co *>* 1 causes faster constriction and leads to a different shape of poles. Previous attempts at modeling division of *E. coli* agree, that a sufficient initial Z-ring force is necessary for the start of the constriction[15, 16].

The subsequent progress of the division is then driven by three time scales. First, by the characteristic production time scale, which we define by the ratio of the active zone volume *V*_a_ and the volume rate Ω

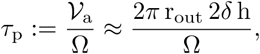

where h := r_out_*−*r_in_ is the current thickness of the active zone and r_out_, r_in_ is the current outer and inner radius respectively, the approximation being valid in the constriction phase. Every *τ*_p_ the sPG grows by the volume *V*_a_. Second, the characteristic remodeling time is defined by the driving force f_Z_ and the solidity modulus *η*, related via the characteristic length 2*δ*, the width of the active zone,

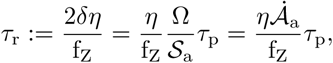

where *S*_a_ is the area of the of the active zone’s front and

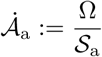

is the arc length rate, i.e. the speed by which the growing PG elongates along the inner membrane. The remodeling time scale expresses how fast the material reshapes under loading. The last, third time scale (growth time scale) is given by the ratio of the ‘driving’ pressure P_tur_ and the resistance modulus *κ*

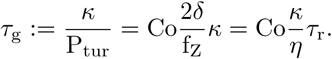

It describes how fast the newly grown sPG distribution diverges due to pressure differences within the thin PG cell wall. Two additional dimensionless numbers can be constructed from these timescales and used to characterize the mode of division in our model. Analyzing the division process using these numbers reveals the deep relation of our quasistatic growth model with the rheology of viscoelastic fluids: The first is an analogue of the Weissenberg number[68, 69]

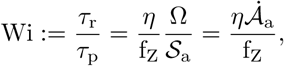

which we define as the ratio of the remodeling and production time scale and a short calculation shows that it is indeed a ratio of ‘viscous’ friction forces to elastic forces: a virtual remodeling force line density *ηA*^.^_a_ (which has the same dimension as a viscous friction force) and the elastic force line density f_Z_. The second dimensionless number is specific to our model and relates the remodeling and growth process by a simple ratio

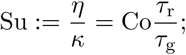

we call Su the susceptibility number as it is a ratio of the pressure susceptibility *κ^−^*^1^ of the PG deposition and the PG fluidity *η^−^*^1^. As these two dimensionless numbers Wi and Su fix only the ratio of the three time scales, the three remodeling and growth parameters *η*, *κ*, and Ω can be scaled appropriately such that the total division time fits the experimental data.

For investigating the model’s sensitivity to the Wi and Su numbers we defined several numerical measures characterising the division process: The wedge formation time (the time instant when the thickness of the PG in the middle is triple the initial thickness, relative to the total division time), the initiation of septation (the time instant when the IM constricts two times faster than the OM, relative to the total division time), the final septum width (relative to the original width of the cell), the mean thickness of the newly grown septal PG (relative to the initial PG thickness), the ratio of the final septum and the length the newly grown sPG measured along the inner membrane (differentiating between constriction and septal division), and the length of the hypotenuse over the reduction in the outer radius and over the elongation of the cell due to division (relative to the length of the arc of the septal PG along the inner membrane).

All theses measures are plotted against Wi and Su in Fig. 4. It is obvious that the main parameter is the Weissenberg number, while the susceptibility number varies the quantities just mildly. At the same time all the variations show the same pattern, meaning only one of them can be selected independently and the others then follow. This indicates, that all mutants lie ”on a line”, between the extrema given in Fig. 4m and t. Hence, our model predicts for example that thick septa go hand in hand with ”flat” constrictions Fig. 4c,f. The one dimensional nature of the change is reflected by the boundaries between the phenotypes indicated in Fig. 4a-f. The fact, that our model is both extremely restricted, and yet predicts several observed phenotypes, which in turn are representative of the morphological variability of divisome mutants can be interpreted as a mechanistic validation.

**Fig. 4.**
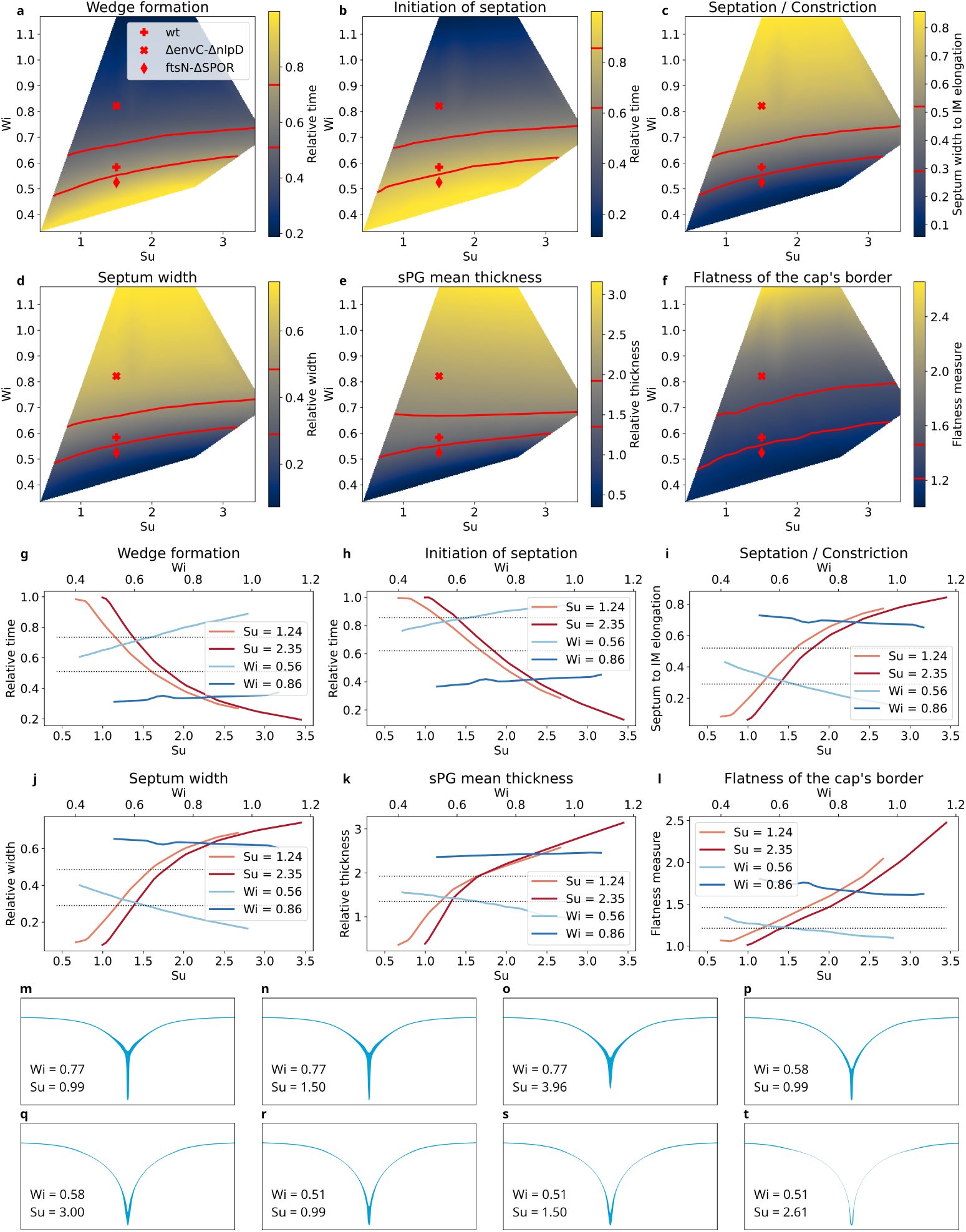
Qualitative analysis of the model’s sensitivity to the Weissenberg (Wi) and Susceptibility (Su) number. **a**-**f**, Contour plots showing selected measures and their dependence on Wi and Su. Points corresponding to the wild type and two mutants marked by red symbols; the three division modes corresponding to them are separated by red isolines. **g**-**l**, Slices through the contour plots from (**a**)-(**f** ) for selected values of Wi or Su, values of selected measures separating the division modes plotted by dotted lines. **m**-**t**, Final morphologies of the division site for selected values of Wi and Su.

### Model Limitations and Outlook

When formulating the present model, we have assumed the material to keep its properties constant during the division. In particular, there is no difference between the old and new septal PG concerning mechanical parameters; the density, stiffness, and anisotropy remain the same. We also have not taken into account potential growth and remodeling rate differences with respect to distance from the inner and outer boundaries and neither have we incorporated the mechanical contribution of the membranes. The lipid layers alone can carry approximately a few percent of the load and bending torque that PG can [18, 70, 71]. Their mechanical contributions hence must be expected to not play a major role, however, increasing the membranes’ stiffness may influence the structure, presumably via bending forces and thereby balance defects in PG caused e.g. by genetic mutations and sustain the division process[72]. As the model accounts only for mass increase and *continuous* deformation, the final cytokinesis phase is out of scope of our current work. To some extent, this is a technicality, as we could perform some infinitesimal ”surgery” to separate bacteria at the end of the cytokinesis stage and to close off septum formation. However, due to its short duration, there is also very little experimental data on the event of cytokinesis, making validation difficult.

## Conclusion

Our model faithfully reproduces the constriction and septation phase of *E. coli* cell division in a quantitative manner and with a minimal number of arbitrarily adjustable parameters. Moreover, all our model parameters have a clear physical interpretation and varying the free parameters reproduces phenotypes of mutated bacteria. Our model maps to a limited scope of phenotypes, this is reflected by the limited variability of phenotypes resulting from divisome mutations. These results indicate that our model captures the essential mechanobiology involved in the constriction and septation phases of cell division. The growth equation is fully coupled with mechanics and coarse-grained, effective enzyme kinetics, it is not prescribed by a separate, artificial evolutionary equation and allows for a variable thickness of the PG. In contrast to other approaches, our model relies on simple thermodynamic assumptions, namely the dependence of chemical potential on stress and a quasi-equilibrium state due to slow growth. It illustrates the universality of stress-mediated growth phenomena in biology. Its conceptual simplicity has allowed us to avoid difficult kinetic attribution questions and extensive parametrization. Our results also highlight the commonality and differences to known models of solid and fluid mechanics. A viscoelastic or elastoplastic material will reduce stress and minimize free energy by accomodating external stress. Growth processes can transfer this type of behavior to ”hard” materials by collocation of active remodeling sites and material incorporation. At the same time, growth leads to a different morphology unknown from inert materials, so that e.g. necking and scission are suppressed in our model. We are optimistic, that the insights gained in this way will transfer to e.g. cocci and perhaps even plant cell division. It remains to be seen whether our model will be able to also treat cytokinesis and the reconstruction of the bacterial poles.

## Methods

### Numerical Modelling

The morphoelastic model was solved using the finite element method. The solver was implemented using FEniCS[73] and uses dynamic remeshing via Gmsh[74]. See the SI for full details about the implementation. The solver will be made publicly available upon acceptance of the paper.

### Image Processing

Images were processed as in[13], the bacterial width was estimated using a square frame of the full images shown in the SI. The model data were then scaled to the width of the image. Note, that we have tested the stability of the model with respect to small variations in the bacterial diameter.

### Cryo-EM specimen preparation

Bacterial strains were grown overnight in LB media, back diluted 1 : 1000 and incub-ated shaking at 37*^◦^*C, 250 rpm to OD_600_ = 0.3. Cells were harvested by centrifugation (2 min, 5000*×*g, RT) and resuspended in LB media to a final OD_600_ = 0.6. This cell suspension (3 *µ*l) was applied to Cflat-2/1 200 mesh copper or gold grids (Electron Microscopy Sciences) that were glow discharged for 30 seconds at 15 mA. Grids were plunge-frozen in liquid ethane[75] with a FEI Vitrobot Mark IV (Thermo Fisher Scientific) at RT, 100% humidity with a waiting time of 13 seconds, one-side blotting time of 13 seconds and blotting force of 10. Customized parafilm sheets were used for one-side blotting. All subsequent grid handling and transfers were performed in liquid nitrogen. Grids were clipped onto cryo-FIB autogrids (Thermo Fisher Scientific).

### Cryo-FIB milling

Grids were loaded in an Aquilos 2 Cryo-FIB (Thermo Fisher Scientific). Specimen was sputter coated inside the cryo-FIB chamber with inorganic platinum, and an integrated gas injection system (GIS) was used to deposit an organometallic platinum layer to protect the specimen surface and avoid uneven thinning of cells. Cryo-FIB milling was performed at a nominal tilt angle of 14*^◦^ −* 18*^◦^* which translates into a milling angle of 7*^◦^ −* 11*^◦^*[76]. Cryo-FIB milling was performed in several steps of decreasing ion beam currents ranging from 0.5 nA to 10 pA and decreasing thickness to obtain 150*−*250 nm lamellae.

### Cryo-TEM data acquisition

All imaging was done on a FEI Titan Krios (Thermo Fisher Scientific) transmission electron microscope operated at 300 KeV equipped with a Gatan BioQuantum K3 energy filter (20 eV zero-loss filtering) and a Gatan K3 direct electron detector. Prior to data acquisition, a full K3 gain reference was acquired, and ZLP and BioQuantum energy filter were finely tuned. Micrographs were collected with a nominal defocus of -3.5 *µ*m. Data collection was performed in the nanoprobe mode using the SerialEM[77] or Thermo Scientific Tomography 6 software.

## Authors’ contributions

PP conducted research and wrote the FEM model implementation. PP and CA designed research. PP and CA wrote the paper. CA supervised and initiated the project. PPN contributed cryo-electron micrographs. PPN, AV and LC revised the paper and contributed to the biological interpretation of the results.

## Acknowledgements

P.P. and C.A. were funded by GAUK PRIMUS Grant PRIMUS/20/SCI/015. P.P.N. was funded by a Swiss National Science Foundation (SNSF) Starting Grant TMSGI3 218251. L.H.C. was funded by the National Institutes of Health (R35GM142553). This work has been supported by Charles University Research Centre program No. UNCE/24/SCI/005. C. A. also thanks Mark Dostaĺık for discussions and contributions to early work.

## Footnotes

^1^The letter M represents the Mandel stress tensor, the superscript d indicates the deviatoric part

## Notes

### Competing Interest Statement

The authors have declared no competing interest.

