## Supporting Information for "Stress-mediated growth determines *E. coli* division site morphogenesis"

September 11, 2024

### 1 Theory of elastic growth

#### 1.1 Kinematics

Let $\vec{\chi}$ denote the deformation map which transforms the initial, reference configuration $\mathcal{B} \subset \mathbb{R}^3$ (not necessarily unstressed) to the current configuration $\mathcal{B}_t \subset \mathbb{R}^3$

$$\vec{\chi}\colon \mathcal{B} \to \mathcal{B}_t, \qquad \vec{\chi}\colon \vec{X} \mapsto \vec{x}, \tag{1}$$

where we use the notation $\vec{X}$ and $\vec{x}$ for the position vectors representing points in the reference and current configurations, respectively; the displacement field is defined as

$$\vec{U} := \vec{x} - \vec{X}. \tag{2}$$

The deformation differential is defined as

$$\mathbb{F} := \frac{\partial \vec{\chi}}{\partial \vec{X}} = \nabla \vec{\chi}, \tag{3}$$

maps material vectors, tangent to the reference configuration, on spatial vectors, tagent to the current configurationGurtin et al. [2010], and can be represented by a $3 \times 3$ matrix. Note that

$$\mathbb{F} = \mathbb{I} + \nabla \vec{U}. \tag{4}$$

In accordance with the theory of morphoelasticity Rodriguez et al. [1994], Goriely [2017], we introduce the elastic tensor $\mathbb{A}$ and the growth tensor $\mathbb{G}$ by the multiplicative split

$$\mathbb{F} := \mathbb{A}\mathbb{G}, \tag{5}$$

where the growth tensor $\mathbb{G}$ first maps the material vectors on the so-called structural vectors, which are then mapped by the elastic tensor $\mathbb{A}$ on the spatial vectors; the composition $\mathbb{A}\mathbb{G}$ may be represented by multiplication of two $3 \times 3$ matricesGurtin et al. [2010]. The so called elastic part $\mathbb{A}$ of the overall, observed local deformation $\mathbb{F}$ compensates for the residual stress induced by the growth and in effect maintains the material integrity and its mechanical equilibrium. The growth tensor $\mathbb{G}$ describes both the local change of shape due to structural changes and insertion of new material, which increases the local density. The relation between the Lagrangian density $\rho_{\mathrm{R}}$, Eulerian density $\rho$, and the structural density $\rho_{\mathrm{s}}$ is

$$\rho_{\mathrm{R}} := \rho \det \mathbb{F} = \rho \det \mathbb{A} \, \det \mathbb{G} =: \rho_{\mathrm{s}} \det \mathbb{G}, \tag{6}$$

where det denotes determinant of a linear mapping Gurtin et al. [2010]. We assume the material properties of the new, septal PG are the same as of the original one, hence the structural density $\rho_{\mathrm{s}}$ is treated as a material's constant.

In particular, in the notation of Goriely [2017] $\dot{\rho}_{\mathrm{s}} = 0$. For the total mass $\mathcal{M}$, total volume $\mathcal{V}$, and their rates we have the following relations

$$\mathcal{M} = \int_{\mathcal{B}} \rho_{\mathrm{R}} \mathrm{d}X, \quad \dot{\mathcal{M}} = \int_{\mathcal{B}} \dot{\rho}_{\mathrm{R}}\, \mathrm{d}X, \quad \mathcal{V} := \frac{\mathcal{M}}{\rho_{\mathrm{s}}}, \quad \dot{\mathcal{V}} = \int_{\mathcal{B}} \dot{\overline{(\det \mathbb{G})}}\, \mathrm{d}X. \tag{7}$$

Under these assumptions the precise value of $\rho_{\mathrm{s}}$ need not to be specified in the model as it completely factors out from the equations. From experimental date we determine the total volume rate rather than the total mass rate and the energy density, see the next section, is measured in experiments also per volume, not per mass.

#### 1.2 Elasticity

We consider elastic energy of the so called simple material Truesdell and Noll [2004], Gurtin et al. [2010]

$$\mathcal{F}(\vec{\chi}, \mathbb{G}) = \int_{\mathcal{B}} \rho_{\mathrm{s}} \psi(\nabla \vec{\chi} \mathbb{G}^{-1})(\det \mathbb{G})\, \mathrm{d}X, \tag{8}$$

where $\psi$ denotes the so-called energy storage function, have units $\mathrm{J} \cdot \mathrm{kg}^{-1}$, and depends solely on the elastic strain $\mathbb{A} = \nabla \vec{\chi} \mathbb{G}^{-1}$ at a particular point $\vec{X}$ in an algebraic way; here $\mathbb{G}^{-1}$ denotes the inverse linear mapping to $\mathbb{G}$, which can be also represented by a square matrix. Since the cell is in a mechanical equilibrium, the differential of the elastic energy $\mathcal{F}$ with respect to the deformation, denoted $\mathrm{D}_{\vec{\chi}} \mathcal{F}$, has to equal the virtual power of the external loading Gurtin et al. [2010]. Denoting the virtual Lagrangian velocity field by $\vec{V}$ and the outer normal vector on $\partial \mathcal{B}$ by $\vec{n}_R$, the variation of the energy can be rewritten by a chain rule

$$\begin{aligned}
\langle \mathrm{D}_{\vec{\chi}} \mathcal{F}, \vec{V} \rangle &= \lim_{\varepsilon \to 0} \frac{1}{\varepsilon} \Big( \mathcal{F}(\mathbb{F}(\vec{\chi} + \varepsilon \vec{V}), \mathbb{G}) - \mathcal{F}(\mathbb{F}(\vec{\chi}), \mathbb{G}) \Big) \\
&= \int_{\mathcal{B}} \frac{\partial(\rho_{\mathrm{s}} \psi)}{\partial \mathbb{A}} : \nabla \vec{V} \mathbb{G}^{-1} (\det \mathbb{G})\, \mathrm{d}X \\
&= -\int_{\mathcal{B}} \mathrm{div} \left( \frac{\partial(\rho_{\mathrm{s}} \psi)}{\partial \mathbb{A}} \mathbb{G}^{-\top} (\det \mathbb{G}) \right) \cdot \vec{V}\, \mathrm{d}X \\
&\quad + \int_{\partial \mathcal{B}} \left( \frac{\partial(\rho_{\mathrm{s}} \psi)}{\partial \mathbb{A}} \mathbb{G}^{-\top} (\det \mathbb{G}) \vec{n}_R \right) \cdot \vec{V}\, \mathrm{d}S,
\end{aligned}$$

where : stands for contraction over two adjacent indices, $\cdot$ over one index, and div denotes divergence, here of a tensor fieldGurtin et al. [2010]. Denoting the external bulk force $\vec{b}$ and the surface force $\vec{\ell}$, the virtual power balance reads

$$\int_{\mathcal{B}} \mathbb{P} : \nabla \vec{V}\, \mathrm{d}X = \int_{\mathcal{B}} \vec{b} \cdot \vec{V}\, \mathrm{d}X + \int_{\partial \mathcal{B}} \vec{\ell} \cdot \vec{V}\, \mathrm{d}S, \tag{9}$$

with the first Piola–Krchoff stress tensor $\mathbb{P}$ and Kirchhoff stress tensor[1] $\mathbb{K}$

$$\mathbb{P} := \frac{\partial(\rho_{\mathrm{s}}\psi)}{\partial\mathbb{A}}\mathbb{G}^{-\top}(\det\mathbb{G}) = \mathbb{K}\mathbb{F}^{-\top}(\det\mathbb{G}),$$
$$\mathbb{K} := \frac{\partial(\rho_{\mathrm{s}}\psi)}{\partial\mathbb{A}}\mathbb{A}^{\top}.$$

By localizing the power balance one arrives at the so called strong or classical, point-wise formulation Gurtin et al. [2010]

$$-\operatorname{div}\mathbb{P} = \vec{b}, \quad \text{in } \mathcal{B},$$
$$\mathbb{P}\vec{n}_R = \vec{\ell}, \quad \text{on } \partial\mathcal{B}.$$

Regarding the loading, we assume that $\partial\mathcal{B} = \partial\mathcal{B}_{\mathrm{out}} \cup \partial\mathcal{B}_{\mathrm{in}}$ where $\partial\mathcal{B}_{\mathrm{out}}$ denotes the outer shell of the capsule and $\partial\mathcal{B}_{\mathrm{in}}$ denotes the inner shell of the capsule. The pressure load $\vec{\ell}_\pi$ is then

$$\vec{\ell}_\pi = -\pi\,(\operatorname{Cof}\mathbb{F})\,\vec{n}_{\mathrm{R}} \quad \text{on } \partial\mathcal{B}, \tag{10}$$

where Cof denotes the cofactor matrixGurtin et al. [2010], $\pi \in \mathbb{R}$ is the difference between the in-cell pressure and the outer atmospheric pressure, that is

$$\pi = 0 \quad \text{on } \partial\mathcal{B}_{\mathrm{out}},$$
$$\pi = \mathrm{P}_{\mathrm{tur}} \quad \text{on } \partial\mathcal{B}_{\mathrm{in}},$$

with $\mathrm{P}_{\mathrm{tur}}$ standing for the turgor pressure. The Z-ring load is given by

$$\vec{\ell}_{\mathrm{Z}} = -\mathrm{P}_{\mathrm{Z}}\zeta(a)|(\operatorname{Cof}\mathbb{F})\vec{n}_{\mathrm{R}}|\vec{e}_{\hat{R}} \quad \text{on } \partial\mathcal{B}_{\mathrm{in}}, \tag{11}$$

where $a$ denotes the arc length distance from the midplane, $\vec{e}_{\hat{R}}$ is the basis vector in the radial direction, $\mathrm{P}_{\mathrm{Z}} = \mathrm{F}_{\mathrm{Z}}/S$ is the area-averaged Z-ring force with the weight

$$S = \int_{\partial\mathcal{B}_{\mathrm{in}}} \zeta(a)|(\operatorname{Cof}\mathbb{F})\vec{n}_{\mathrm{R}}|\,\mathrm{d}S,$$

and $\zeta : (-\infty, +\infty) \to [0, 1]$ is the localization function

$$\zeta(a) = \exp\left(1 - \frac{\delta^4}{(a-\delta)^2(a+\delta)^2}\right) \tag{12}$$

within the zring radius $\delta > 0$ and 0 otherwise. Putting it all together the virtual power balance reads

$$\int_{\mathcal{B}} \mathbb{P} : \nabla\vec{V}\,\mathrm{d}X = -\int_{\partial\mathcal{B}_{\mathrm{in}}} \left(\mathrm{P}_{\mathrm{tur}}\,(\operatorname{Cof}\mathbb{F})\,\vec{n}_{\mathrm{R}} + \mathrm{P}_{\mathrm{Z}}\zeta(a)|(\operatorname{Cof}\mathbb{F})\vec{n}_{\mathrm{R}}|\vec{e}_{\hat{R}}\right)\cdot\vec{V}\,\mathrm{d}S, \tag{13}$$

[1] Unlike in the theory of plasticity, here $\det\mathbb{G} \neq 1$; however, we prefer to keep $\det\mathbb{G}$ together with the differential form $\mathrm{d}X$ and define the Kirchhoff stress tensor without it.

for all virtual Lagrangian velocity fields $\vec{V}$.

To close the system of equations for the deformation field $\vec{\chi}$, we prescribe a specific form of the free energy $\psi$. We split the elastic tensor $\mathbb{A}$ into volumetric part $J$ and isochoric part $\bar{\mathbb{A}}$ defined respectively

$$J := \det \mathbb{A}, \qquad \bar{\mathbb{A}} := J^{-\frac{1}{3}}\mathbb{A};$$

note that upon linearization, with $\mathbb{A} = \mathbb{I} + \mathbb{H}$, $|\mathbb{H}| \to 0$, the splitting turns into the trace $\mathrm{tr}(\mathbb{A})$ and deviatoric part $\mathbb{A}^{\mathrm{d}}$

$$J = 1 + \mathrm{tr}(\mathbb{A}) + \mathrm{O}(|\mathbb{H}|^2), \qquad \bar{\mathbb{A}} = \mathbb{I} + \mathbb{A}^{\mathrm{d}} + \mathrm{O}(|\mathbb{H}|^2), \quad \mathbb{A}^{\mathrm{d}} := \mathbb{A} - \frac{1}{3}(\mathrm{tr}\,\mathbb{A})\mathbb{I} = \mathbb{H}^{\mathrm{d}}.$$

Using this splitting, we consider the following ansatz for the free energy[2]

$$\rho_{\mathrm{s}}\psi(\mathbb{A}) = K\psi_{\mathrm{v}}(J) + \mu\psi_{\mathrm{i}}(\bar{\mathbb{A}}) + \beta\psi_{\mathrm{ani}}(\mathbb{A}), \tag{14}$$

where $K$ is the bulk modulus, $\mu$ is the shear modulus, and $\beta$ is the anisotropic modulus. The particular contributions then read

$$\psi_{\mathrm{v}}(J) = \frac{1}{50}\big(J^5 + J^{-5} - 2\big), \qquad \psi_{\mathrm{i}}(\bar{\mathbb{A}}) = \frac{1}{54}\left(|\bar{\mathbb{A}}|^6 - 27\right),$$

and

$$\psi_{\mathrm{ani}}(\mathbb{A}) = |\mathbb{A}\vec{m}_{\mathrm{R}}|^2 - 1 - \ln|\mathbb{A}\vec{m}_{\mathrm{R}}|^2, \tag{15}$$

where $\vec{m}_{\mathrm{R}}$ denotes a unit vector in the reference configuration that represents the prevalent fiber orientation. Specifically, we consider circumferential pattern, i.e. $\vec{m}_{\mathrm{R}} = \vec{e}_{\Theta}$. The anisotropic energy term $\psi_{\mathrm{ani}}$ has the same quadratic envelope as another widely considered formula

$$\tilde{\psi}(\mathbb{A}) = \frac{1}{2}\left(|\mathbb{A}\vec{m}_{\mathrm{R}}|^2 - 1\right)^2,$$

which is referred to as the *standard fiber-reinforcing model*, see e.g. Demirkoparan and Pence [2008], Merodio and Ogden [2002], Horgan and Murphy [2011].

This choice of energy complies with all the mathematical requirements such as polyconvexity and thanks to the isochoric splitting the first two coefficients, $K$ and $\mu$, have direct mechanical interpretation.

The derivative of the free energy density then reads

$$\begin{aligned}\frac{\partial(\rho_{\mathrm{s}}\psi)}{\partial\mathbb{A}} &= \left(K\frac{\partial\psi_{\mathrm{v}}}{\partial J}J\mathbb{I} + \left(\mu\frac{\partial\psi_{\mathrm{i}}}{\partial\bar{\mathbb{A}}}\bar{\mathbb{A}}^{\top}\right)^{\mathrm{d}}\right)\mathbb{A}^{-\top} + \beta\frac{\partial\psi_{\mathrm{ani}}}{\partial\mathbb{A}}\\ &= \mathbb{A}^{-\top}\left(K\frac{\partial\psi_{\mathrm{v}}}{\partial J}J\mathbb{I} + \left(\mu\bar{\mathbb{A}}^{\top}\frac{\partial\psi_{\mathrm{i}}}{\partial\bar{\mathbb{A}}}\right)^{\mathrm{d}}\right) + \beta\frac{\partial\psi_{\mathrm{ani}}}{\partial\mathbb{A}},\end{aligned}$$

[2]The structural density $\rho_{\mathrm{s}}$ itself is used for conceptual reasons only, but it is otherwise completely irrelevant in our model; hence it is not specified. The values of the moduli $K$, $\mu$, $\beta$ determine the elastic response.

where

$$\frac{\partial \psi_{\rm v}}{\partial J} = \frac{1}{10}\left(J^4 - J^{-6}\right), \quad \frac{\partial \psi_{\rm i}}{\partial \bar{\mathbb{A}}} = \frac{1}{9}|\bar{\mathbb{A}}|^4\bar{\mathbb{A}}, \quad \frac{\partial \psi_{\rm ani}}{\partial \mathbb{A}} = 2\left(1 - \frac{1}{|\mathbb{A}\vec{m}_{\rm R}|^2}\right)\mathbb{A}\vec{m}_{\rm R}\otimes\vec{m}_{\rm R}.$$

The Kirchhoff stress tensor then equals

$$\mathbb{K} = K\frac{\partial \psi_{\rm v}}{\partial J}J\mathbb{I} + \left(\mu\frac{\partial \psi_{\rm i}}{\partial \bar{\mathbb{A}}}\bar{\mathbb{A}}^\top\right)^{\rm d} + \beta\frac{\partial \psi_{\rm ani}}{\partial \mathbb{A}}\mathbb{A}^\top.$$

The quadratic envelope of $\psi$ has the following engineering moduli (see the Appendix 4.2 for a detailed derivation)

$$\mathrm{E}_g = \frac{9K\mu}{3K+\mu} + 4\beta, \quad \mathrm{E}_p = \mu\frac{9K\mu + 12K\beta + 4\mu\beta}{\mu^2 + 3K\mu + 3K\beta + 4\mu\beta}, \quad \nu_{gp} = \frac{1}{2}\frac{3K - 2\mu}{3K+\mu}.$$

For

$$K = 172.00\,\mathrm{MPa}, \qquad \mu = 6.96\,\mathrm{MPa}, \qquad \beta = 5.36\,\mathrm{MPa},$$

the Young moduli and Poisson ratio equals the desired values

$$\mathrm{E}_g \approx 42\,\mathrm{MPa}, \qquad \mathrm{E}_p \approx 23\,\mathrm{MPa}, \qquad \nu_{gp} \approx 0.48.$$

#### 1.3 Growth and remodeling

As we assume the transformation law to be mechanically driven, we will seek a relation between the rate $\dot{\mathbb{G}}$ and the driving force $\Xi := -\mathrm{D}_{\mathbb{G}}\mathcal{F}$. Recalling the free energy (8) we observe

$$\Xi := -\mathrm{D}_{\mathbb{G}}\mathcal{F} = \mathbb{A}^\top\frac{\partial(\rho_{\rm s}\psi)}{\partial \mathbb{A}}\mathbb{G}^{-\top}(\det\mathbb{G}) - (\rho_{\rm s}\psi)\mathbb{G}^{-\top}(\det\mathbb{G}).$$

The relation between the rate $\dot{\mathbb{G}}$ and the driving force $\Xi$ may be written via a convex dissipation potential $\mathcal{R}(\vec{\chi},\mathbb{G},\dot{\mathbb{G}})$ and its Legendre conjugate with respect to the last variable $\mathcal{R}^*(\vec{\chi},\mathbb{G},\Xi)$, either as a force balance or as a rate equationMielke and Roubíček [2015], Roubíček [2022]

$$-\mathrm{D}_{\mathbb{G}}\mathcal{F} = \mathrm{D}_{\dot{\mathbb{G}}}\mathcal{R}(\vec{\chi},\mathbb{G},a,\dot{\mathbb{G}}) \quad\Longleftrightarrow\quad \dot{\mathbb{G}} = \mathrm{D}_{\Xi}\mathcal{R}^*(\vec{\chi},\mathbb{G},a,-\mathrm{D}_{\mathbb{G}}\mathcal{F}). \tag{16}$$

The second assumption on the transformation law is that the total mass rate is constant during the time, in principle determined by the cell's production capacity of precursors. Since

$$\dot{\overline{(\det\mathbb{G})}} = (\det\mathbb{G})\mathbb{G}^{-\top}\colon\dot{\mathbb{G}} = \mathrm{tr}(\dot{\mathbb{G}}\mathbb{G}^{-1})\det\mathbb{G}, \tag{17}$$

the total volume increase is given by

$$\dot{\mathcal{V}} = \int_{\mathcal{B}}\dot{\overline{(\det\mathbb{G})}}\,\mathrm{d}X = \int_{\mathcal{B}}\mathrm{tr}(\dot{\mathbb{G}}\mathbb{G}^{-1})(\det\mathbb{G})\,\mathrm{d}X = \int_{\mathcal{B}}\mathrm{tr}(\mathbb{L}_\kappa)(\det\mathbb{G})\,\mathrm{d}X. \tag{18}$$

We consider the dissipation potential of the form

$$\mathcal{R}(\vec{\chi}, \mathbb{G}, a, \dot{\mathbb{G}}) = \begin{cases} \int_{\mathcal{B}} \left( \frac{1}{\zeta(a)} \mathrm{R}(\mathbb{L}_\kappa) - (\rho_\mathrm{s}\psi) \operatorname{tr}(\mathbb{L}_\kappa) \right) (\det \mathbb{G}) \, \mathrm{d}X, & \text{if } \dot{\mathcal{V}} = \Omega \\ +\infty, & \text{otherwise,} \end{cases},$$

where R is the dissipation potential density and $\Omega$ is the prescribed global volume rate[3], since the transformation law then relates the transformation rate $\mathbb{L}_\kappa$ with the driving force given by the Mandel stress[4] $\mathbb{M}$

$$\mathbb{L}_\kappa := \dot{\mathbb{G}}\mathbb{G}^{-1}, \qquad \mathbb{M} := \mathbb{A}^\top \frac{\partial(\rho_\mathrm{s}\psi)}{\partial \mathbb{A}} = -\mathrm{D}_\mathbb{G}\mathcal{F}\mathbb{G}^\top (\det \mathbb{G})^{-1} + (\rho_\mathrm{s}\psi)\mathbb{I}, \tag{19}$$

as it is also common in the theory of plasticityGurtin et al. [2010]. The variation of $\mathcal{R}$ in the direction $\tilde{\mathbb{V}}$ reads

$$\langle \mathrm{D}_{\dot{\mathbb{G}}} \mathcal{R}(\vec{\chi}, \mathbb{G}, a, \dot{\mathbb{G}}), \tilde{\mathbb{V}} \rangle = \int_{\mathcal{B}} \left( \frac{1}{\zeta(a)} \frac{\partial \mathrm{R}}{\partial \mathbb{L}_\kappa} - (\rho_\mathrm{s}\psi)\mathbb{I} - \omega \mathbb{I} \right) : \tilde{\mathbb{V}}\mathbb{G}^{-1} (\det \mathbb{G}) \, \mathrm{d}X,$$

where $\omega$ is the Lagrange multiplier stemming from the global volume rate constraint, and hence the transformation law (16) becomes (when testing the equation with $\tilde{\mathbb{V}} = \mathbb{V}\mathbb{G}$)

$$\int_{\mathcal{B}} \left( \frac{\partial \mathrm{R}}{\partial \mathbb{L}_\kappa} - \omega \zeta(a) \mathbb{I} \right) : \mathbb{V} (\det \mathbb{G}) \, \mathrm{d}X = \int_{\mathcal{B}} \zeta(a) \mathbb{M} : \mathbb{V} (\det \mathbb{G}) \, \mathrm{d}X,$$
$$\int_{\mathcal{B}} \operatorname{tr}(\dot{\mathbb{G}}\mathbb{G}^{-1}) (\det \mathbb{G}) \, \mathrm{d}X = \Omega,$$

with the Mandel stress $\mathbb{M}$ given by the free energy ansatz (14)

$$\mathbb{M} = K \frac{\partial \psi_\mathrm{v}}{\partial J} J \mathbb{I} + \left( \mu \bar{\mathbb{A}}^\top \frac{\partial \psi_\mathrm{i}}{\partial \bar{\mathbb{A}}} \right)^\mathrm{d} + \beta \mathbb{A}^\top \frac{\partial \psi_\mathrm{ani}}{\partial \mathbb{A}}.$$

Following Goriely [2017], we split the transformation rate $\dot{\mathbb{G}}\mathbb{G}^{-1}$ into the growth rate $\gamma$ and the remodeling rate $\Gamma$ defined by

$$\gamma := \mathrm{l}_\kappa := \operatorname{tr}(\mathbb{L}_\kappa) = \operatorname{tr}(\dot{\mathbb{G}}\mathbb{G}^{-1}) = \dot{g} g^{-1},$$
$$\Gamma := \mathbb{L}_\kappa^\mathrm{d} := \mathbb{L}_\kappa - \frac{1}{3} (\operatorname{tr} \mathbb{L}_\kappa) \mathbb{I} = (\dot{\mathbb{G}}\mathbb{G}^{-1})^\mathrm{d} = \dot{\bar{\mathbb{G}}} \bar{\mathbb{G}}^{-1},$$

where

$$g := \det \mathbb{G}, \qquad \bar{\mathbb{G}} := g^{-\frac{1}{3}} \mathbb{G},$$

[3]Note that $\rho_\mathrm{s}\psi$ is given by (14) in terms of elastic moduli, where $\rho_\mathrm{s}$ disappears.

[4]From the same reasons as for the Kirchhoff stress, we define the Mandel stress without $\det \mathbb{G}$.

are the volumetric and isochoric part of $\mathbb{G}$ respectively. The strong formulations of the growth law, remodeling law and the volume rate constraint read

$$
\begin{aligned}
\frac{\partial \mathrm{R}}{\partial \mathrm{l}_\kappa} - \omega\zeta(a) &= \frac{\zeta(a)}{3}\operatorname{tr}(\mathbb{M}), && \text{in } \mathcal{B},\\
\frac{\partial \mathrm{R}}{\partial \mathbb{L}_\kappa^{\mathrm{d}}} &= \zeta(a)\mathbb{M}^{\mathrm{d}}, && \text{in } \mathcal{B},\\
\int_{\mathcal{B}} \operatorname{tr}(\dot{\mathbb{G}}\mathbb{G}^{-1})(\det \mathbb{G})\,\mathrm{d}X &= \Omega, && \text{in } \mathbb{R},
\end{aligned}
$$

where

$$
\begin{aligned}
\operatorname{tr}(\mathbb{M}) &= 3K\frac{\partial \psi_{\mathrm{v}}}{\partial J}J + \beta\frac{\partial \psi_{\mathrm{ani}}}{\partial \mathbb{A}} : \mathbb{A},\\
\mathbb{M}^{\mathrm{d}} &= \left(\mu\bar{\mathbb{A}}^\top\frac{\partial \psi_{\mathrm{i}}}{\partial \bar{\mathbb{A}}}\right)^{\mathrm{d}} + \beta\left(\mathbb{A}^\top\frac{\partial \psi_{\mathrm{ani}}}{\partial \mathbb{A}}\right)^{\mathrm{d}}.
\end{aligned}
$$

As linear relations between the Mandel stress $\mathbb{M}$ and transformation rate $\mathbb{L}_\kappa$ is sufficient, we consider quadratic R

$$
\mathrm{R}(\mathbb{L}_\kappa) = \frac{1}{2}\mathbb{L}_\kappa : \mathcal{D}\mathbb{L}_\kappa,
$$

where $(\mathcal{D})_{ijkl} = \mathrm{D}_{ijkl}$ is a symmetric, positive definite four order tensor. As there is no reason to suppose the contrary, we assume the law is isotropic and hence the whole tensor $\mathcal{D}$ reduces to two moduli, the resistance $\kappa$ and solidity $\eta$, i.e.

$$
\mathrm{R}(\mathbb{L}_\kappa) = \kappa\mathrm{R}_{\mathrm{v}}(\operatorname{tr}\mathbb{L}_\kappa) + \eta\mathrm{R}_{\mathrm{i}}(\mathbb{L}_\kappa^{\mathrm{d}}) = \frac{\kappa}{2}\operatorname{tr}^2(\mathbb{L}_\kappa) + \eta|\mathbb{L}_\kappa^{\mathrm{d}}|^2,
$$

with

$$
\frac{\partial \mathrm{R}}{\partial \mathbb{L}_\kappa} = \kappa\frac{\partial \mathrm{R}_{\mathrm{v}}}{\partial \mathrm{l}_\kappa}\mathbb{I} + \eta\frac{\partial \mathrm{R}_{\mathrm{i}}}{\partial \mathbb{L}_\kappa^{\mathrm{d}}} = \kappa\operatorname{tr}(\mathbb{L}_\kappa)\mathbb{I} + 2\eta\mathbb{L}_\kappa^{\mathrm{d}}.
$$

For $\kappa = \frac{2}{3}\eta$ the relation simplifies to $\mathrm{R}(\mathbb{L}_\kappa) = \eta|\mathbb{L}_\kappa|^2$.

Since the hydrostatic pressure $p$ is one third of the trace of the Cauchy stress

$$
\mathbb{T} := (\det \mathbb{A})^{-1}\frac{\partial(\rho_{\mathrm{s}}\psi)}{\partial \mathbb{A}}\mathbb{A}^\top = -p\mathbb{I} + \mathbb{T}^{\mathrm{d}},
$$

we have

$$
\begin{aligned}
\mathbb{M} &= (\det \mathbb{A})\mathbb{A}^\top(\det \mathbb{A})^{-1}\frac{\partial(\rho_{\mathrm{s}}\psi)}{\partial \mathbb{A}}\mathbb{A}^\top\mathbb{A}^{-\top} = (\det \mathbb{A})\mathbb{A}^\top\mathbb{T}\mathbb{A}^{-\top},\\
\frac{1}{3}\operatorname{tr}(\mathbb{M}) &= -\frac{1}{3}\operatorname{tr}(\mathbb{T})(\det \mathbb{A}) = -p(\det \mathbb{A}),
\end{aligned}
$$

and the growth law therefore depends on the hydrostatic pressure $p$ via

$$\frac{\partial \mathrm{R}}{\partial \mathrm{l}_\kappa} = \kappa \mathrm{l}_\kappa = \kappa\gamma = \kappa\,\mathrm{tr}(\mathbb{L}_\kappa) = \kappa \dot{g} g^{-1} = \zeta(a)\left((-p)(\det \mathbb{A}) + \omega\right).$$

For the volume rate we have then

$$\dot{\mathcal{V}} = \int_{\mathcal{B}} \dot{g} g^{-1} (\det \mathbb{G})\,\mathrm{d}X = \int_{\mathcal{B}} \zeta(a)\left(-\frac{P}{\kappa} + \gamma_0\right)(\det \mathbb{G})\,\mathrm{d}X,$$

where we defined the pressure field $P$ and referential growth rate $\gamma_0$ by

$$P := p(\det \mathbb{A}), \qquad \gamma_0 := \frac{\omega}{\kappa} = \frac{P_0}{\kappa},$$

i.e. the Lagrange multiplier $\omega$ plays the role of the referential pressure $P_0$. The phenomenological growth law depending on pressure is justified by thermodynamic considerations in the following sub-subsection.

##### 1.3.1 Thermodynamic justification

We consider the PG layer as an open system, with the actual PG a space-filling background, and the number of precursors $N$ fixed and in equilibrium. The spherical stress in the system is represented by an inhomogeneous pressure $p = -\operatorname{tr}\mathbb{T}$. The Maxwell relation

$$\left(\frac{\partial \mu_i}{\partial p}\right)_{\mu_{j\neq i}, V, T} = \left(\frac{\partial V}{\partial n_i}\right)_{\mu, T} = V_{m,i} \tag{20}$$

is readily derived from the grand canonical potential. $V_{m,i}$ is the partial molar volume of species $i$. We express the total differential of the chemical potential of the PG precursor as

$$\mathrm{d}\mu_{PG}(p, n_{PG}, T) = \left(\frac{\partial \mu_{PG}}{\partial p}\right)\mathrm{d}p + \left(\frac{\partial \mu_{PG}}{\partial T}\right)\mathrm{d}T + \left(\frac{\partial \mu_{PG}}{\partial n_{PG}}\right)\mathrm{d}n_{PG}. \tag{21}$$

Where we assume ideality in the sense that

$$\left(\frac{\partial \mu_i}{\partial n_j}\right)_{j\neq i, p, T} = 0. \tag{22}$$

and

$$\left(\frac{\partial \mu_{PG}}{\partial n_{PG}}\right)_{p, T, n\neq PG} = RT\frac{1}{\chi_{PG}}. \tag{23}$$

Here, we have introduced the precursor mole fraction

$$\chi_{PG} = \frac{\rho_{PG}}{\sum_i \rho_i}, \tag{24}$$

which depends on its own and all other mass densities $\rho_i$. In our PG shell, the pressure and precursor density is a function of the position $\mathbf{r}$, but temperature is constant. Hence,

$$\mu_{PG}(\mathbf{r}) = \mu^0_{PG} + \int_{p^0}^{p(\mathbf{r})} V_{m,PG} \mathrm{d}p + RT \ln \frac{\chi(\mathbf{r})_{PG}}{\chi^0_{PG}}. \tag{25}$$

Here the superscript $^0$ indicates a reference value. We will assume furthermore, that the partial molar volume of PG does not depend on pressure, so that the integral reduces to

$$[p(\mathbf{r}) - p^0]V_{m,PG}. \tag{26}$$

We assume the system is in equilibrium, hence the chemical potential in all portions of the system is the same:

$$\mu_{PG}(\mathbf{r}) = \mu_{PG}(\mathbf{r}'). \tag{27}$$

Plugging this into our equation reduces to

$$p(\mathbf{r})V_{m,PG} + RT \ln \rho(\mathbf{r})_{PG} = p(\mathbf{r}')V_{m,PG} + RT \ln \rho(\mathbf{r}')_{PG}. \tag{28}$$

Or, choosing a reference state somewhere in the system:

$$\rho(\mathbf{r}) = \rho^0 e^{-\beta V_{m,PG}[p(\mathbf{r})-p^0]}. \tag{29}$$

If the exponent is small, i.e. due to small pressure differences, a Taylor expansion up to first order in the exponent linearizes the equation to

$$\rho(\mathbf{r}) = \rho^0 - \rho^0 \beta V_{m,PG}[p(\mathbf{r}) - p^0]. \tag{30}$$

This makes the concentration linearly dependent on pressure. The growth kinetics can then be assumed to be of the form

$$\dot{g}g^{-1} \propto \rho(\mathbf{r}) \propto -(p(\mathbf{r}) - p^0). \tag{31}$$

#### 1.4 Eikonal equation

The signed distance function $\hat{a} : \mathcal{B}_t \to \mathbb{R}$, measuring on the deformed cell wall $\mathcal{B}_t$ the distance from the symmetry midplane of the bacterium, satisfies the so-called Eikonal equationGallot et al. [2004]

$$\begin{aligned} |\nabla_{\vec{x}} \hat{a}| &= 1, \qquad \text{in } \mathcal{B}_t, \\ \hat{a} &= 0, \qquad \text{on the midplane of } \mathcal{B}_t, \end{aligned}$$

where $\nabla_{\vec{x}} \hat{a} = \frac{\partial a}{\partial \vec{x}}$ is the differential with respect to the current position $\vec{x}$. For the pulled back signed distance function $a : \mathcal{B} \to \mathbb{R}$, $a(\vec{X}) := \hat{a}(\vec{x})$ for $\vec{x} = \vec{\chi}(\vec{X})$, we use the chain rule $\frac{\partial a}{\partial \vec{x}} = \mathbb{F}^{-\top}\nabla a$, with $\nabla a = \frac{\partial a}{\partial \vec{X}}$, to transform the differential equation into the reference configuration $\mathcal{B}$

$$\begin{aligned} |\mathbb{F}^{-\top}\nabla a| = |\nabla a|_{\mathbb{C}^{-1}} = 1, &\qquad \text{in } \mathcal{B}, \\ a = 0, &\qquad \text{on the midplane of } \mathcal{B}, \end{aligned}$$

where $\mathbb{C} := \mathbb{F}^\top\mathbb{F}$ is the right Cauchy–Green tensor and

$$|\vec{v}|^2_{\mathbb{C}^{-1}} = (\vec{v}, \vec{v})_{\mathbb{C}^{-1}} = \vec{v} \cdot \mathbb{C}^{-1}\vec{v},$$

for any co-vector $\vec{v}$.

### 2 Numerical implementation

We discretize the problem in space and time and use arbitrary Lagrangian-Eulerian method (ALE) with remeshing.

#### 2.1 Mixed finite element formulation

For spatial discretization of the deformation field $\vec{\chi}$, growth tensor $\mathbb{G}$, and the arc-length distance $a$ we use finite element method (FEM)Babuška and Strouboulis [2001], Ciarlet [2002]. In order to overcome numerical instabilities we also enrich the finite element space for the deformation $\vec{\chi}$ via the so-called Hu-Washizu variational principleSimo et al. [1985], Lamichhane [2009], Faghih Shojaei and Yavari [2019]. We introduce new variables, which help in approximating the nonlinearities

$$\iota \approx J = \det \mathbb{A}, \qquad -p \approx \frac{1}{\mu} \left.\frac{\partial(\rho_{\mathrm{s}}\psi)}{\partial J}\right|_{J=\iota},$$

and the auxiliary Lagrangian

$$\mathcal{L}(\vec{\chi}, \iota, p, \mathbb{G}) = \frac{1}{\mu}\int_{\mathcal{B}} \rho_{\mathrm{s}}\psi\left(\iota^{\frac{1}{3}}\bar{\mathbb{A}}\right)(\det \mathbb{G})\,\mathrm{d}X - \int_{\mathcal{B}} p(\det \mathbb{A} - \iota)(\det \mathbb{G})\,\mathrm{d}X,$$

where

$$\mathbb{A} = (\nabla\vec{\chi})\mathbb{G}^{-1}, \quad \bar{\mathbb{A}} = \overline{(\nabla\vec{\chi})}\bar{\mathbb{G}}^{-1}, \quad \det \mathbb{A} = \det(\nabla\vec{\chi})(\det \mathbb{G})^{-1}.$$

The unknown fields are then sought in the following spaces

$$\begin{aligned}
\vec{\chi} \in \mathbf{W}_h \quad &:= \{\vec{V} \in \mathrm{C}(\mathcal{B}; \mathbb{R}^3) \quad &: \vec{V}|_K \in \mathbf{P}_2(K), \forall K \in \mathcal{T}_h\}, \\
\iota \in V_h \quad &:= \{\varphi \in \mathrm{C}(\mathcal{B}) \quad &: \varphi|_K \in P_1(K), \forall K \in \mathcal{T}_h\}, \\
p \in V_h \quad &:= \{\varphi \in \mathrm{C}(\mathcal{B}) \quad &: \varphi|_K \in P_1(K), \forall K \in \mathcal{T}_h\}, \\
\mathbb{G} \in \mathbb{V}_h \quad &:= \{\mathbb{V} \in \mathrm{C}(\mathcal{B}; \mathbb{R}^{3\times 3}) \quad &: \mathbb{V}|_K \in \mathbb{P}_1(K), \forall K \in \mathcal{T}_h\}, \\
a \in W_h \quad &:= \{\varphi \in \mathrm{C}(\mathcal{B}) \quad &: \varphi|_K \in P_2(K), \forall K \in \mathcal{T}_h\},
\end{aligned}$$

where normal font denotes spaces of scalar functions, bold of vector functions, and black board of $\mathbb{R}^{3\times 3}$-tensor functions, C denotes the space of continuous functions, and $\mathcal{T}_h$ is the triangular mesh with elements $K$. We use Gauss quadrature of degree 8.

##### 2.1.1 Elasticity

The mixed formulation is then obtained by variation of the auxiliary Lagrangian $\mathcal{L}$ that is balanced by the virtual work of the pressure and Z-ring load

$$
\begin{aligned}
\langle \mathrm{D}_{\vec{X}}\mathcal{L}, \vec{V}\rangle &= -\int_{\partial\mathcal{B}_{\mathrm{in}}} \left( \frac{\mathrm{P}_{\mathrm{tur}}}{\mu} \left(\operatorname{Cof}\mathbb{F}\right) \vec{n}_{\mathrm{R}} + \frac{\mathrm{P}_{\mathrm{Z}}}{\mu}\zeta(a)|(\operatorname{Cof}\mathbb{F})\vec{n}_{\mathrm{R}}|\vec{e}_{\hat{R}} \right) \cdot \vec{V}\,\mathrm{d}S, \quad && \forall \vec{V} \in \mathbf{W}_h,\\
\langle \mathrm{D}_p\mathcal{L}, q\rangle &= 0, && \forall q \in V_h,\\
\langle \mathrm{D}_\iota\mathcal{L}, \upsilon\rangle &= 0, && \forall \upsilon \in V_h,
\end{aligned}
$$

and the particular variations read

$$
\begin{aligned}
\langle \mathrm{D}_{\vec{X}}\mathcal{L}, \vec{V}\rangle &= \int_{\mathcal{B}} \frac{1}{\mu}\mathbb{P} : \vec{V}\,\mathrm{d}X,\\
\langle \mathrm{D}_p\mathcal{L}, q\rangle &= -\int_{\mathcal{B}} (\det\mathbb{A} - \iota) q (\det\mathbb{G})\,\mathrm{d}X,\\
\langle \mathrm{D}_\iota\mathcal{L}, \upsilon\rangle &= \int_{\mathcal{B}} \left( \frac{\iota^{-1}}{3}\left[\frac{1}{\mu}\operatorname{tr}(\mathbb{K})\right]_{\mathbb{A}=\iota^{\frac{1}{3}}\bar{\mathbb{A}}} + p \right) \upsilon(\det\mathbb{G})\,\mathrm{d}X,
\end{aligned}
$$

where

$$
\frac{1}{\mu}\mathbb{P} = \frac{1}{\mu}\mathbb{K}\mathbb{F}^{-\top}(\det\mathbb{G}),
$$

and for the purpose of this discrete mixed formulation we define

$$
\frac{1}{\mu}\mathbb{K} := -p(\det\mathbb{A})\mathbb{I} + \left[\frac{1}{\mu}\mathbb{K}^{\mathrm{d}}\right]_{\mathbb{A}=\iota^{\frac{1}{3}}\bar{\mathbb{A}}},
$$

$$
\frac{1}{\mu}\mathbb{K}^{\mathrm{d}} := \left(\frac{\partial\psi_{\mathrm{i}}}{\partial\bar{\mathbb{A}}}\bar{\mathbb{A}}^{\top}\right)^{\mathrm{d}} + \frac{\beta}{\mu}\left(\frac{\partial\psi_{\mathrm{ani}}}{\partial\mathbb{A}}\mathbb{A}^{\top}\right)^{\mathrm{d}}, \tag{32}
$$

$$
\frac{1}{\mu}\operatorname{tr}(\mathbb{K}) := 3\frac{K}{\mu}\frac{\partial\psi_{\mathrm{v}}}{\partial J}J + \frac{\beta}{\mu}\frac{\partial\psi_{\mathrm{ani}}}{\partial\mathbb{A}} : \mathbb{A}. \tag{33}
$$

##### 2.1.2 Growth and remodeling

In the transformation law (16) we replace the driving force by

$$
\Xi := -\mu\mathrm{D}_{\mathbb{G}}\mathcal{L}.
$$

The weak formulation after dividing by the sear modulus $\mu$ then reads

$$
\begin{aligned}
\int_{\mathcal{B}} \left(\frac{1}{\mu}\frac{\partial\mathrm{R}}{\partial\mathbb{L}_\kappa} - \omega\zeta(a)\mathbb{I}\right) : \mathbb{V}(\det\mathbb{G})\,\mathrm{d}X &= \int_{\mathcal{B}} \zeta(a)\frac{1}{\mu}\mathbb{M} : \mathbb{V}(\det\mathbb{G})\,\mathrm{d}X, \quad \forall \mathbb{V} \in \mathbb{V}_h\\
\int_{\mathcal{B}} \operatorname{tr}(\dot{\mathbb{G}}\mathbb{G}^{-1})(\det\mathbb{G})\,\mathrm{d}X &= \Omega,
\end{aligned}
$$

where for the purpose of this discrete mixed formulation we define

$$\frac{1}{\mu}\mathbb{M} := -p\iota\mathbb{I} + \left[\frac{1}{\mu}\mathbb{M}^{\mathrm{d}}\right]_{\mathbb{A}=\iota^{\frac{1}{3}}\bar{\mathbb{A}}},$$
$$\frac{1}{\mu}\mathbb{M}^{\mathrm{d}} := \left(\bar{\mathbb{A}}^{\top}\frac{\partial\psi_{\mathrm{i}}}{\partial\bar{\mathbb{A}}}\right)^{\mathrm{d}} + \frac{\beta}{\mu}\left(\mathbb{A}^{\top}\frac{\partial\psi_{\mathrm{ani}}}{\partial\mathbb{A}}\right)^{\mathrm{d}}. \tag{34}$$

##### 2.1.3 Arc-length

The equation for arc-length is regularized by an elliptic term in which the prefactor $\varepsilon$ is chosen appropriately. At the initial time we find the solution by decreasing $\varepsilon = 1$ to $\varepsilon = 0.04 \cdot h_{max}$, where $h_{max}$ is the diameter of the largest mesh element. This minimal value of $\varepsilon$ is then kept for later time steps. The equation then reads

$$\varepsilon\int_{\mathcal{B}}\mathbb{C}^{-1}\nabla a : \nabla\varphi\,\mathrm{d}X + \int_{\mathcal{B}}|\mathbb{F}^{-\top}\nabla a|\varphi\,\mathrm{d}X = \int_{\mathcal{B}}\varphi\,\mathrm{d}X, \quad \forall\varphi\in V_h$$
$$a = 0, \qquad \text{on the midplane of } \mathcal{B}.$$

#### 2.2 Time discretization

We use a fractional time stepping scheme, where in the half step we solve first the equations for $(\vec{\chi}, \iota, p, \mathbb{G})$ by a monolithic solver and then in the full step solve the equation for the arc-length $a$; we iterate this until the relative difference between the two subsequent steps is smaller than a given tolerance, here $10^{-4}$.

For solving the $(\vec{\chi}, \iota, p, \mathbb{G})$-system we use the minimizing movements time discretizationAmbrosio [1995], which gives for the equations of elasticity

$$\int_{\mathcal{B}}\frac{1}{\mu}\mathbb{P}_{n+\frac{1}{2}} : \nabla\vec{V}\,\mathrm{d}X$$
$$= -\int_{\partial\mathcal{B}_{\mathrm{in}}}\left(\frac{\mathrm{P}_{\mathrm{tur}}}{\mu}\left(\mathrm{Cof}\,\mathbb{F}_{n+\frac{1}{2}}\right)\vec{n}_{\mathrm{R}} + \frac{\mathrm{P}_{\mathrm{Z}}}{\mu}\zeta(a_n)|(\mathrm{Cof}\,\mathbb{F}_{n+\frac{1}{2}})\vec{n}_{\mathrm{R}}|\vec{e}_{\hat{R}}\right)\cdot\vec{V}\,\mathrm{d}S, \tag{35}$$

$$0 = \int_{\mathcal{B}}(\det\mathbb{A}_{n+\frac{1}{2}} - \iota_{n+\frac{1}{2}})(\det\mathbb{G}_{n+\frac{1}{2}})q\,\mathrm{d}X, \tag{36}$$

$$0 = \int_{\mathcal{B}}\left(\frac{\iota^{-1}_{n+\frac{1}{2}}}{3}\left[\frac{1}{\mu}\mathrm{tr}(\mathbb{K})\right]_{\mathbb{A}=(\iota_{n+\frac{1}{2}})^{\frac{1}{3}}\bar{\mathbb{A}}_{n+\frac{1}{2}}} + p_{n+\frac{1}{2}}\right)(\det\mathbb{G}_{n+\frac{1}{2}})\upsilon\,\mathrm{d}X, \tag{37}$$

where

$$\frac{1}{\mu}\mathbb{P}_{n+\frac{1}{2}} = \frac{1}{\mu}\mathbb{K}_{n+\frac{1}{2}}\mathbb{F}^{-\top}_{n+\frac{1}{2}}(\det\mathbb{G}_{n+\frac{1}{2}}),$$

and we define

$$\frac{1}{\mu}\mathbb{K}_{n+\frac{1}{2}} := -p_{n+\frac{1}{2}}(\det\mathbb{A}_{n+\frac{1}{2}})\mathbb{I} + \left[\frac{1}{\mu}\mathbb{K}^{\mathrm{d}}\right]_{\mathbb{A}=(\iota_{n+\frac{1}{2}})^{\frac{1}{3}}\bar{\mathbb{A}}_{n+\frac{1}{2}}},$$

while $\frac{1}{\mu}\mathbb{K}^{\mathrm{d}}$ with $\frac{1}{\mu}\operatorname{tr}(\mathbb{K})$ are given by the constitutive equations (32) and (33). For the transformation law the minimizing movement discretization yields

$$\begin{aligned}\int_{\mathcal{B}} \left(\frac{1}{\mu}\frac{\partial \mathrm{R}}{\partial \mathbb{L}_\kappa}\left((\mathbb{L}_\kappa)_{n+\frac{1}{2}}\right) - \omega_{n+\frac{1}{2}}\zeta(a_n)\mathbb{I}\right) &: \mathbb{V}(\det \mathbb{G}_n)\,\mathrm{d}X \\ = \int_{\mathcal{B}} \zeta(a_n)\frac{1}{\mu}\mathbb{M}_{n+\frac{1}{2}}\mathbb{G}^{-\top}_{n+\frac{1}{2}}\mathbb{G}^{\top}_n &: \mathbb{V}(\det \mathbb{G}_{n+\frac{1}{2}})\,\mathrm{d}X, \quad (38) \\ \Omega = \int_{\mathcal{B}} \operatorname{tr}\left((\mathbb{L}_\kappa)_{n+\frac{1}{2}}\right)(\det \mathbb{G}_n)\,\mathrm{d}X, & \quad (39)\end{aligned}$$

with

$$\begin{aligned}(\mathbb{L}_\kappa)_{n+\frac{1}{2}} &:= \frac{\mathbb{G}_{n+\frac{1}{2}} - \mathbb{G}_n}{\tau}\mathbb{G}_n^{-1}, \\ \frac{1}{\mu}\mathbb{M}_{n+\frac{1}{2}} &:= -p_{n+\frac{1}{2}}\iota_{n+\frac{1}{2}}\mathbb{I} + \left[\frac{1}{\mu}\mathbb{M}^{\mathrm{d}}\right]_{\mathbb{A}=(\iota_{n+\frac{1}{2}})^{\frac{1}{3}}\bar{\mathbb{A}}_{n+\frac{1}{2}}},\end{aligned}$$

where $\frac{1}{\mu}\mathbb{M}^{\mathrm{d}}$ is given by the constitutive relation (34). With computed $\vec{\chi}_{n+\frac{1}{2}}$ and $\mathbb{G}_{n+\frac{1}{2}}$ we then solve the equation for $a_{n+1}$

$$\begin{aligned}\varepsilon\int_{\mathcal{B}} \mathbb{C}^{-1}_{n+\frac{1}{2}}\nabla a_{n+1} : \nabla\varphi\,\mathrm{d}X + \int_{\mathcal{B}} |\mathbb{F}^{-\top}_{n+\frac{1}{2}}\nabla a_{n+1}|\varphi\,\mathrm{d}X &= \int_{\mathcal{B}} \varphi\,\mathrm{d}X, \\ a_{n+1} &= 0, \qquad \text{on the midplane of } \mathcal{B},\end{aligned}$$

where $\mathbb{C}_{n+\frac{1}{2}} = \mathbb{F}^{\top}_{n+\frac{1}{2}}\mathbb{F}_{n+\frac{1}{2}}$.

#### 2.3 ALE method

The non-uniform growth degenerates the elasticity equation and hence the tensor of elastic 'constants' becomes ill-conditioned; we therefore want to filter the growth out. After several ALE moves the mesh degenerates and we remesh the domain completely.

We want to find a global approximation of the natural, stress-free configuration, which is given by ALE displacement $\vec{\phi} : \mathbb{R}^3 \to \mathbb{R}^3$

$$\vec{\phi} \in \operatorname{ArgMin} \frac{1}{2}\int_{\mathcal{B}} |(\mathbb{I} + \nabla\vec{\phi})\mathbb{G}^{-1} - \mathbb{I}|^2(\det \mathbb{G})\,\mathrm{d}X.$$

Given the solution $\vec{\phi}$ we define the ALE deformation

$$\vec{\chi}_{\mathrm{ALE}}(\vec{X}) := \vec{X} + \vec{\phi}, \quad \vec{\chi}_{\mathrm{ALE}} : \mathcal{B} \to \tilde{\mathcal{B}}, \quad \tilde{\mathcal{B}} := \vec{\chi}_{\mathrm{ALE}}(\mathcal{B}),$$

The displacement and growth tensor are then updated

$$\begin{aligned}&\tilde{\vec{U}} : \mathbb{R}^3 \to \mathbb{R}^3, \quad \tilde{\vec{U}} := \vec{U} - \vec{\phi}, \quad \tilde{\vec{\chi}} : \tilde{\mathcal{B}} \to \mathcal{B}_t, \quad \tilde{\vec{\chi}}(\tilde{X}) := \vec{\chi}({\vec{\chi}_{\mathrm{ALE}}}^{-1}(\tilde{X})), \\ &\tilde{\mathbb{F}} := \nabla\tilde{\vec{\chi}} = \mathbb{F}(\nabla\vec{\chi}_{\mathrm{ALE}})^{-1}, \quad \tilde{\mathbb{G}} := \mathbb{G}(\nabla\vec{\chi}_{\mathrm{ALE}})^{-1},\end{aligned}$$

while the elastic tensor and growth rate remain unchanged

$$\tilde{\mathbb{A}} := \tilde{\mathbb{F}}\tilde{\mathbb{G}}^{-1} = \nabla\tilde{\vec{\chi}}(\nabla\vec{\chi}_{\mathrm{ALE}})\mathbb{G}^{-1} = \mathbb{F}(\nabla\vec{\chi}_{\mathrm{ALE}})^{-1}\tilde{\mathbb{G}}^{-1} = \nabla\vec{\chi}\mathbb{G}^{-1} = \mathbb{A},$$
$$\tilde{\mathbb{L}}_{\kappa} := \dot{\tilde{\mathbb{G}}}\tilde{\mathbb{G}}^{-1} = \dot{\mathbb{G}}(\nabla\vec{\chi}_{\mathrm{ALE}})^{-1}(\nabla\vec{\chi}_{\mathrm{ALE}})\mathbb{G}^{-1} = \mathbb{L}_{\kappa}.$$

The Lagrangian transforms as

$$\begin{aligned}\mathcal{L}(\vec{\chi}, \mathbb{G}, \iota, p) &= \frac{1}{\mu}\int_{\tilde{\mathcal{B}}} \rho_{\mathrm{s}}\psi\left(\iota^{\frac{1}{3}}\bar{\mathbb{A}}\right)(\det\mathbb{G})(\det\nabla\vec{\chi}_{\mathrm{ALE}})^{-1}\,\mathrm{d}\tilde{X} \\ &\quad - \int_{\tilde{\mathcal{B}}} p(\det\mathbb{A} - \iota)(\det\mathbb{G})(\det\nabla\vec{\chi}_{\mathrm{ALE}})^{-1}\,\mathrm{d}\tilde{X} \\ &= \frac{1}{\mu}\int_{\tilde{\mathcal{B}}} \rho_{\mathrm{s}}\psi\left(\iota^{\frac{1}{3}}\bar{\tilde{\mathbb{A}}}\right)(\det\tilde{\mathbb{G}})\,\mathrm{d}\tilde{X} - \int_{\tilde{\mathcal{B}}} p(\det\tilde{\mathbb{A}} - \iota)(\det\tilde{\mathbb{G}})\,\mathrm{d}\tilde{X} \\ &= \tilde{\mathcal{L}}(\tilde{\vec{\chi}}, \tilde{\mathbb{G}}, \iota, p).\end{aligned}$$

The loading transforms as

$$\begin{aligned}&\int_{\partial\mathcal{B}_{\mathrm{in}}} \left(\frac{\mathrm{P}_{\mathrm{tur}}}{\mu}(\mathrm{Cof}\,\mathbb{F})\,\vec{n}_{\mathrm{R}} + \frac{\mathrm{P}_{\mathrm{Z}}}{\mu}\zeta(a)|(\mathrm{Cof}\,\mathbb{F})\vec{n}_{\mathrm{R}}|\vec{e}_{\hat{R}}\right)\cdot\vec{V}\,\mathrm{d}S \\ &= \int_{\partial\tilde{\mathcal{B}}_{\mathrm{in}}} \left(\frac{\mathrm{P}_{\mathrm{tur}}}{\mu}(\mathrm{Cof}\,\mathbb{F})\,(\mathrm{Cof}\,\nabla\tilde{\vec{\chi}})^{-1}\tilde{\vec{n}}_{\mathrm{R}} + \frac{\mathrm{P}_{\mathrm{Z}}}{\mu}\zeta(a)|(\mathrm{Cof}\,\mathbb{F})(\mathrm{Cof}\,\nabla\tilde{\vec{\chi}})^{-1}\tilde{\vec{n}}_{\mathrm{R}}|\vec{e}_{\hat{R}}\right)\cdot\vec{V}\,\mathrm{d}\tilde{S} \\ &= \int_{\partial\tilde{\mathcal{B}}_{\mathrm{in}}} \left(\frac{\mathrm{P}_{\mathrm{tur}}}{\mu}(\mathrm{Cof}\,\tilde{\mathbb{F}})\tilde{\vec{n}}_{\mathrm{R}} + \frac{\mathrm{P}_{\mathrm{Z}}}{\mu}\zeta(a)|(\mathrm{Cof}\,\tilde{\mathbb{F}})\tilde{\vec{n}}_{\mathrm{R}}|\vec{e}_{\hat{R}}\right)\cdot\vec{V}\,\mathrm{d}\tilde{S}\end{aligned}$$

The total volume rate transforms as

$$\dot{\mathcal{V}} = \int_{\mathcal{B}} \mathrm{tr}(\mathbb{L}_{\kappa})(\det\mathbb{G})\,\mathrm{d}X = \int_{\tilde{\mathcal{B}}} \mathrm{tr}(\mathbb{L}_{\kappa})(\det\mathbb{G})(\det\nabla\vec{\chi}_{\mathrm{ALE}})^{-1}\,\mathrm{d}\tilde{X} = \dot{\tilde{\mathcal{V}}},$$

and the dissipation potential transforms as

$$\begin{aligned}\mathcal{R}(\vec{\chi}, \mathbb{G}, a, \dot{\mathbb{G}}) &= \begin{cases}\displaystyle\int_{\tilde{\mathcal{B}}} \left(\frac{1}{\zeta(a)}\mathrm{R}(\mathbb{L}_{\kappa}) - (\rho_{\mathrm{s}}\psi)\,\mathrm{tr}(\mathbb{L}_{\kappa})\right)(\det\mathbb{G})(\det\nabla\vec{\chi}_{\mathrm{ALE}})^{-1}\,\mathrm{d}\tilde{X}, & \text{if } \dot{\tilde{\mathcal{V}}} = \Omega \\ +\infty, & \text{otherwise,}\end{cases} \\ &= \tilde{\mathcal{R}}(\tilde{\vec{\chi}}, \tilde{\mathbb{G}}, a, \dot{\tilde{\mathbb{G}}}).\end{aligned}$$

### 3 Elastic growth of bacterial wall

The peptidoglycan wall of the rod-like cell is modelled as a hollow capsule with length $L$ and radii of the inner and outer spherical caps denoted by $R_{\mathrm{in}}$ and $R_{\mathrm{out}}$, respectively; see Fig. 1 for details. At the division site, the FtsZ-ring applies a localized and radially inward pointing traction force. Moreover, internal turgor pressure acts from the inside of the cell.

Given the geometric symmetries we reformulate the problem in cylindrical coordinates. In both, material and spatial descriptions, we work with cylindrical

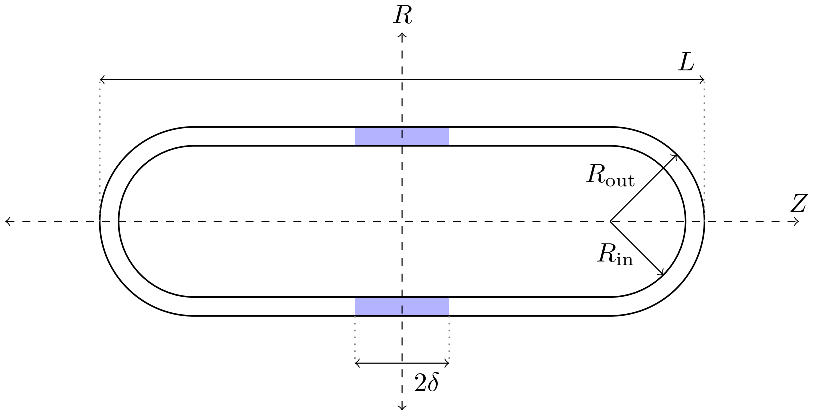

Figure 1: Cell geometry.

coordinates $(R, \Theta, Z)$ and $(r, \theta, z)$, respectively. Although they are identical, we denote them differently in order to distinguish between the reference and current, deformed configuration.

##### 3.1 Plane strain formulation

Given the Lagrangian basis $(\vec{e}_{\hat{R}}, \vec{e}_{\hat{\Theta}}, \vec{e}_{\hat{Z}})$ and its analogue $(\vec{e}_{\hat{r}}, \vec{e}_{\hat{\theta}}, \vec{e}_{\hat{z}})$ in the Eulerian configuration, we assume that the traction forces acting on the bacterial wall have radial orientation and the displacement field is thus axially symmetric, i.e. we consider the following ansatz for the displacement field

$$\vec{U} = \begin{bmatrix} \mathrm{U}^{\hat{r}}(R, Z) \\ 0 \\ \mathrm{U}^{\hat{z}}(R, Z) \end{bmatrix}. \tag{40}$$

In cylindrical coordinates the deformation gradient reads

$$\mathbb{F} = \mathbb{I} + \nabla\vec{U} = \begin{bmatrix} 1 + \frac{\partial \mathrm{U}^{\hat{r}}}{\partial R} & \frac{1}{R}\frac{\partial \mathrm{U}^{\hat{r}}}{\partial \Theta} - \frac{\mathrm{U}^{\hat{\theta}}}{R} & \frac{\partial \mathrm{U}^{\hat{r}}}{\partial Z} \\ \frac{\partial \mathrm{U}^{\hat{\theta}}}{\partial R} & 1 + \frac{1}{R}\frac{\partial \mathrm{U}^{\hat{\theta}}}{\partial \Theta} + \frac{\mathrm{U}^{\hat{r}}}{R} & \frac{\partial \mathrm{U}^{\hat{\theta}}}{\partial Z} \\ \frac{\partial \mathrm{U}^{\hat{z}}}{\partial R} & \frac{1}{R}\frac{\partial \mathrm{U}^{\hat{z}}}{\partial \Theta} & 1 + \frac{\partial \mathrm{U}^{\hat{z}}}{\partial Z} \end{bmatrix}. \tag{41}$$

However, by virtue of ansatz (40), the formula reduces to

$$\mathbb{F} = \begin{bmatrix} \mathrm{F}^{\hat{r}}{}_{\hat{R}} & 0 & \mathrm{F}^{\hat{r}}{}_{\hat{Z}} \\ 0 & \mathrm{F}^{\hat{\theta}}{}_{\hat{\Theta}} & 0 \\ \mathrm{F}^{\hat{z}}{}_{\hat{R}} & 0 & \mathrm{F}^{\hat{z}}{}_{\hat{Z}} \end{bmatrix} = \begin{bmatrix} 1 + \frac{\partial \mathrm{U}^{\hat{r}}}{\partial R} & 0 & \frac{\partial \mathrm{U}^{\hat{r}}}{\partial Z} \\ 0 & 1 + \frac{\mathrm{U}^{\hat{r}}}{R} & 0 \\ \frac{\partial \mathrm{U}^{\hat{z}}}{\partial R} & 0 & 1 + \frac{\partial \mathrm{U}^{\hat{z}}}{\partial Z} \end{bmatrix}. \tag{42}$$

In the same spirit we postulate the elastic tensor $\mathbb{A}$ has the same structure, although it is in principle not a gradient of a vector field

$$\mathbb{A} = \begin{bmatrix} \mathrm{A}^{\hat{r}}{}_{\hat{r}} & 0 & \mathrm{A}^{\hat{r}}{}_{\hat{z}} \\ 0 & \mathrm{A}^{\hat{\theta}}{}_{\hat{\theta}} & 0 \\ \mathrm{A}^{\hat{z}}{}_{\hat{r}} & 0 & \mathrm{A}^{\hat{z}}{}_{\hat{z}} \end{bmatrix}, \tag{43}$$

where non of the component depends on $\theta$. As the elastic deformation is supposed to be smaller than the growth, we denote the basis in the intermediate, structural configuration also $(\vec{e}_{\hat{r}}, \vec{e}_{\hat{\theta}}, \vec{e}_{\hat{z}})$, just to reduce the complexity of notation. It follows then the growth tensor must have the very same structure

$$\mathbb{G} = \begin{bmatrix} \mathrm{G}^{\hat{r}}{}_{\hat{R}} & 0 & \mathrm{G}^{\hat{r}}{}_{\hat{Z}} \\ 0 & \mathrm{G}^{\hat{\theta}}{}_{\hat{\Theta}} & 0 \\ \mathrm{G}^{\hat{z}}{}_{\hat{R}} & 0 & \mathrm{G}^{\hat{z}}{}_{\hat{Z}} \end{bmatrix}, \tag{44}$$

where also no component is $\Theta$ dependent. We also suppose that in the intermediate configuration, presumed by the multiplicative split (5), both basis $(\vec{e}_{\hat{R}}, \vec{e}_{\hat{\Theta}}, \vec{e}_{\hat{Z}})$ and $(\vec{e}_{\hat{r}}, \vec{e}_{\hat{\theta}}, \vec{e}_{\hat{z}})$ are available and hence we express $\mathbb{G}$ with respect to them. We may assume equivalently that the growth also complies with the cylindrical symmetry. As a consequence, the whole problem is cylindrical symmetric.

The first Piola, Cauchy, Mandel, and Kirchhoff stress has to follow the same structure

$$\mathbb{P} = \begin{bmatrix} \mathrm{P}^{\hat{r}\hat{R}} & 0 & \mathrm{P}^{\hat{r}\hat{Z}} \\ 0 & \mathrm{P}^{\hat{\theta}\hat{\Theta}} & 0 \\ \mathrm{P}^{\hat{z}\hat{R}} & 0 & \mathrm{P}^{\hat{z}\hat{Z}} \end{bmatrix}, \qquad \mathbb{T} = \begin{bmatrix} \mathrm{T}^{\hat{r}\hat{r}} & 0 & \mathrm{T}^{\hat{r}\hat{z}} \\ & \mathrm{T}^{\hat{\theta}\hat{\theta}} & 0 \\ & & \mathrm{T}^{\hat{z}\hat{z}} \end{bmatrix}, \tag{45}$$

$$\mathbb{M} = \begin{bmatrix} \mathrm{M}^{\hat{r}\hat{r}} & 0 & \mathrm{M}^{\hat{r}\hat{z}} \\ 0 & \mathrm{M}^{\hat{\theta}\hat{\theta}} & 0 \\ \mathrm{M}^{\hat{z}\hat{r}} & 0 & \mathrm{M}^{\hat{z}\hat{z}} \end{bmatrix}, \qquad \mathbb{K} = \begin{bmatrix} \mathrm{K}^{\hat{r}\hat{r}} & 0 & \mathrm{K}^{\hat{r}\hat{z}} \\ & \mathrm{K}^{\hat{\theta}\hat{\theta}} & 0 \\ & & \mathrm{K}^{\hat{z}\hat{z}} \end{bmatrix}. \tag{46}$$

We define the two-dimensional counterparts

$$\mathbb{F}_{\mathrm{2D}} := \begin{bmatrix} \mathrm{F}^{\hat{r}}{}_{\hat{R}} & \mathrm{F}^{\hat{r}}{}_{\hat{Z}} \\ \mathrm{F}^{\hat{z}}{}_{\hat{R}} & \mathrm{F}^{\hat{z}}{}_{\hat{Z}} \end{bmatrix}, \ \mathbb{G}_{\mathrm{2D}} := \begin{bmatrix} \mathrm{G}^{\hat{r}}{}_{\hat{R}} & \mathrm{G}^{\hat{r}}{}_{\hat{Z}} \\ \mathrm{G}^{\hat{z}}{}_{\hat{R}} & \mathrm{G}^{\hat{z}}{}_{\hat{Z}} \end{bmatrix}, \ \mathbb{A}_{\mathrm{2D}} := \begin{bmatrix} \mathrm{A}^{\hat{r}}{}_{\hat{r}} & \mathrm{A}^{\hat{r}}{}_{\hat{z}} \\ \mathrm{A}^{\hat{z}}{}_{\hat{r}} & \mathrm{A}^{\hat{z}}{}_{\hat{z}} \end{bmatrix}, \tag{47a}$$

where $\mathbb{A}_{\mathrm{2D}} = \mathbb{F}_{\mathrm{2D}}\mathbb{G}_{\mathrm{2D}}^{-1}$, and

$$\mathbb{P}_{\mathrm{2D}} := \begin{bmatrix} \mathrm{P}^{\hat{r}\hat{R}} & \mathrm{P}^{\hat{r}\hat{Z}} \\ \mathrm{P}^{\hat{z}\hat{R}} & \mathrm{P}^{\hat{z}\hat{Z}} \end{bmatrix}, \qquad \mathbb{T}_{\mathrm{2D}} := \begin{bmatrix} \mathrm{T}^{\hat{r}\hat{r}} & \mathrm{T}^{\hat{r}\hat{z}} \\ & \mathrm{T}^{\hat{z}\hat{z}} \end{bmatrix}, \tag{48}$$

$$\mathbb{M}_{\mathrm{2D}} := \begin{bmatrix} \mathrm{M}^{\hat{r}\hat{r}} & \mathrm{M}^{\hat{r}\hat{z}} \\ \mathrm{M}^{\hat{z}\hat{r}} & \mathrm{M}^{\hat{z}\hat{z}} \end{bmatrix}, \qquad \mathbb{K}_{\mathrm{2D}} := \begin{bmatrix} \mathrm{K}^{\hat{r}\hat{r}} & \mathrm{K}^{\hat{r}\hat{z}} \\ & \mathrm{K}^{\hat{z}\hat{z}} \end{bmatrix}. \tag{49}$$

The Jacobian of the linear map $\mathbb{G}$ with respect to the basis $(\vec{e}_{\hat{R}}, \vec{e}_{\hat{\Theta}}, \vec{e}_{\hat{Z}})$ and $(\vec{e}_{\hat{r}}, \vec{e}_{\hat{\theta}}, \vec{e}_{\hat{z}})$ is the determinant of the matrix

$$\det \begin{bmatrix} \mathrm{G}^{\hat{r}}{}_{\hat{R}} & 0 & \mathrm{G}^{\hat{r}}{}_{\hat{Z}} \\ 0 & \mathrm{G}^{\hat{\theta}}{}_{\hat{\Theta}} & 0 \\ \mathrm{G}^{\hat{z}}{}_{\hat{R}} & 0 & \mathrm{G}^{\hat{z}}{}_{\hat{Z}} \end{bmatrix},$$

in principle differing from $\det \mathbb{G}$. However, as all the basis vectors are orthogonal and and of unit length, both metric tensors corresponding to them are identities with unit determinant. Hence it holds

$$\det \mathbb{G} = \det \begin{bmatrix} \mathrm{G}^{\hat{r}}{}_{\hat{R}} & 0 & \mathrm{G}^{\hat{r}}{}_{\hat{Z}} \\ 0 & \mathrm{G}^{\hat{\theta}}{}_{\hat{\Theta}} & 0 \\ \mathrm{G}^{\hat{z}}{}_{\hat{R}} & 0 & \mathrm{G}^{\hat{z}}{}_{\hat{Z}} \end{bmatrix} = \mathrm{G}^{\hat{\theta}}{}_{\hat{\Theta}}(\det \mathbb{G}_{\mathrm{2D}}).$$

As the decomposition into a 2D tensor and the $(\hat{\Theta}, \hat{\theta})$-component is closed under transposition, invertion, and multiplication, and addition, we have for the first Piola–Kirchhoff stress tensor

$$\frac{1}{\mu}\mathbb{P}_{\mathrm{2D}} = \frac{1}{\mu}\mathbb{K}_{\mathrm{2D}}\mathbb{F}_{\mathrm{2D}}^{-\top}\mathrm{G}^{\hat{\theta}}{}_{\hat{\Theta}}(\det \mathbb{G}_{\mathrm{2D}}), \tag{50}$$

$$\frac{1}{\mu}\mathrm{P}^{\hat{\theta}\hat{\Theta}} = \frac{1}{\mu}\mathrm{K}^{\hat{\theta}\hat{\theta}}(\mathrm{F}^{\hat{\theta}}{}_{\hat{\Theta}})^{-1}\mathrm{G}^{\hat{\theta}}{}_{\hat{\Theta}}(\det \mathbb{G}_{\mathrm{2D}}). \tag{51}$$

##### 3.1.1 Equations of Mechanical Equilibrium

We shall now reformulate the equations of mechanical equilibrium (35) in the cylindrical plane strain setting. Considering the virtual velocity field compatible with the kinematic constraint (40)

$$\vec{V} = \begin{bmatrix} \mathrm{V}^{\hat{r}}(R, Z) \\ 0 \\ \mathrm{V}^{\hat{z}}(R, Z) \end{bmatrix}, \qquad \vec{V}_{\mathrm{2D}} = \begin{bmatrix} \mathrm{V}^{\hat{r}}(R, Z) \\ \mathrm{V}^{\hat{z}}(R, Z) \end{bmatrix},$$

using the formula for the gradient operator in cylindrical coordinates

$$\nabla\vec{V} = \begin{bmatrix} \frac{\partial \mathrm{V}^{\hat{r}}}{\partial R} & \frac{1}{R}\left(\frac{\partial \mathrm{V}^{\hat{\theta}}}{\partial \Theta} - \mathrm{V}^{\hat{\theta}}\right) & \frac{\partial \mathrm{V}^{\hat{r}}}{\partial Z} \\ \frac{\partial \mathrm{V}^{\hat{\theta}}}{\partial R} & \frac{1}{R}\left(\frac{\partial \mathrm{V}^{\hat{\theta}}}{\partial \Theta} + \mathrm{V}^{\hat{r}}\right) & \frac{\partial \mathrm{V}^{\hat{\theta}}}{\partial Z} \\ \frac{\partial \mathrm{V}^{\hat{z}}}{\partial R} & \frac{1}{R}\frac{\partial \mathrm{V}^{\hat{z}}}{\partial \Theta} & \frac{\partial \mathrm{V}^{\hat{z}}}{\partial Z} \end{bmatrix}, \qquad \nabla\vec{V}_{2\mathrm{D}} = \begin{bmatrix} \frac{\partial \mathrm{V}^{\hat{r}}}{\partial R} & \frac{\partial \mathrm{V}^{\hat{r}}}{\partial Z} \\ \frac{\partial \mathrm{V}^{\hat{z}}}{\partial R} & \frac{\partial \mathrm{V}^{\hat{z}}}{\partial Z} \end{bmatrix},$$

and taking into account the particular structure of the Piola–Kirchhoff stress tensor (45), we can rewrite equation (35) as follows

$$\int_{\mathcal{B}} \left[\frac{1}{\mu}(\mathbb{P}_{2\mathrm{D}})_{n+\frac{1}{2}} : \nabla\vec{V}_{2\mathrm{D}} + \frac{1}{R}\frac{1}{\mu}(\mathrm{P}^{\hat{\theta}\hat{\Theta}})_{n+\frac{1}{2}}\mathrm{V}^{\hat{r}}\right] \mathrm{d}X$$
$$= -\int_{\partial\mathcal{B}_{\mathrm{in}}} \left(\frac{\mathrm{P}_{\mathrm{tur}}}{\mu}\left(\mathrm{Cof}\,\mathbb{F}_{n+\frac{1}{2}}\right)\vec{n}_{\mathrm{R}} + \frac{\mathrm{P}_{\mathrm{Z}}}{\mu}\zeta(a_n)|(\mathrm{Cof}\,\mathbb{F}_{n+\frac{1}{2}})\vec{n}_{\mathrm{R}}|\vec{e}_{\hat{R}}\right)\cdot\vec{V}\,\mathrm{d}S.$$

Since none of the quantities in the previous equation depends on $\Theta$, we can integrate this quantity out and arrive at

$$\int_{B} \left[R\,\frac{1}{\mu}(\mathbb{P}_{2\mathrm{D}})_{n+\frac{1}{2}} : \nabla\vec{V}_{2\mathrm{D}} + \frac{1}{\mu}(\mathrm{P}^{\hat{\theta}\hat{\Theta}})_{n+\frac{1}{2}}\mathrm{V}^{\hat{r}}\right] \mathrm{d}R\,\mathrm{d}Z \tag{52}$$

$$= -\int_{(\partial B)_{\mathrm{in}}} R\left(\frac{\mathrm{P}_{\mathrm{tur}}}{\mu}\left(\mathrm{F}^{\hat{\theta}}{}_{\hat{\Theta}}\,\mathrm{Cof}\,\mathbb{F}_{2\mathrm{D}}\right)_{n+\frac{1}{2}}\vec{n}_{\mathrm{R}} + \frac{\mathrm{P}_{\mathrm{Z}}}{\mu}\zeta(a)|(\mathrm{F}^{\hat{\theta}}{}_{\hat{\Theta}}\,\mathrm{Cof}\,\mathbb{F}_{2\mathrm{D}})_{n+\frac{1}{2}}\vec{n}_{\mathrm{R}}|\vec{e}_{\hat{R}}\right)\cdot\vec{V}_{2\mathrm{D}}\,\mathrm{d}l, \tag{53}$$

$$0 = \int_{B} R\left((\mathrm{A}^{\hat{\theta}}{}_{\hat{\theta}})_{n+\frac{1}{2}}(\det\mathbb{A}_{2\mathrm{D}})_{n+\frac{1}{2}} - \iota_{n+\frac{1}{2}}\right)(\mathrm{G}^{\hat{\theta}}{}_{\hat{\Theta}})_{n+\frac{1}{2}}(\det\mathbb{G}_{2\mathrm{D}})_{n+\frac{1}{2}}\,q\,\mathrm{d}R\,\mathrm{d}Z, \tag{54}$$

$$0 = \int_{B} R\left(\frac{\iota^{-1}_{n+\frac{1}{2}}}{3}\left[\frac{1}{\mu}\,\mathrm{tr}(\mathbb{K})\right]_{\mathbb{A}=(\iota_{n+\frac{1}{2}})^{\frac{1}{3}}\bar{\mathbb{A}}_{n+\frac{1}{2}}} + p_{n+\frac{1}{2}}\right)(\mathrm{G}^{\hat{\theta}}{}_{\hat{\Theta}})_{n+\frac{1}{2}}(\det\mathbb{G}_{2\mathrm{D}})_{n+\frac{1}{2}}\,\upsilon\,\mathrm{d}R\,\mathrm{d}Z, \tag{55}$$

where $B$ denotes a "half-stadium" shown in Figure 2, $\mathrm{P}_{\mathrm{Z}} = \mathrm{F}_{\mathrm{Z}}/S$ is the averaged Z-ring force with the weight

$$S = \int_{(\partial B)_{\mathrm{in}}} R\zeta(a)|(\mathrm{F}^{\hat{\theta}}{}_{\hat{\Theta}}\,\mathrm{Cof}\,\mathbb{F}_{2\mathrm{D}})\vec{n}_{\mathrm{R}}|\,\mathrm{d}S,$$

and the tensors are given by

$$\frac{1}{\mu}(\mathbb{P}_{2\mathrm{D}})_{n+\frac{1}{2}} = \frac{1}{\mu}(\mathbb{K}_{2\mathrm{D}})_{n+\frac{1}{2}}(\mathbb{F}_{2\mathrm{D}}^{-\top})_{n+\frac{1}{2}}(\mathrm{G}^{\hat{\theta}}{}_{\hat{\Theta}})_{n+\frac{1}{2}}(\det\mathbb{G}_{2\mathrm{D}})_{n+\frac{1}{2}},$$
$$\frac{1}{\mu}(\mathrm{P}^{\hat{\theta}\hat{\Theta}})_{n+\frac{1}{2}} = \frac{1}{\mu}(\mathrm{K}^{\hat{\theta}\hat{\theta}})_{n+\frac{1}{2}}(\mathrm{F}^{\hat{\theta}}{}_{\hat{\Theta}})^{-1}_{n+\frac{1}{2}}(\mathrm{G}^{\hat{\theta}}{}_{\hat{\Theta}})_{n+\frac{1}{2}}(\det\mathbb{G}_{2\mathrm{D}})_{n+\frac{1}{2}},$$

where we define

$$\frac{1}{\mu}(\mathbb{K}_{2\mathrm{D}})_{n+\frac{1}{2}} := -p_{n+\frac{1}{2}}(\mathrm{A}^{\hat{\theta}}{}_{\hat{\theta}})_{n+\frac{1}{2}}(\det \mathbb{A}_{2\mathrm{D}})_{n+\frac{1}{2}}\mathbb{I}_{2\mathrm{D}} + \left[\frac{1}{\mu}\mathbb{K}^{\mathrm{d}}_{2\mathrm{D}}\right]_{\mathbb{A}=(\iota_{n+\frac{1}{2}})^{\frac{1}{3}}\bar{\mathbb{A}}_{n+\frac{1}{2}}},$$

$$\frac{1}{\mu}(\mathrm{K}^{\hat{\theta}\hat{\theta}})_{n+\frac{1}{2}} := -p_{n+\frac{1}{2}}(\mathrm{A}^{\hat{\theta}}{}_{\hat{\theta}})_{n+\frac{1}{2}}(\det \mathbb{A}_{2\mathrm{D}})_{n+\frac{1}{2}} + \left[\frac{1}{\mu}(\mathbb{K}^{\mathrm{d}})^{\hat{\theta}\hat{\theta}}\right]_{\mathbb{A}=(\iota_{n+\frac{1}{2}})^{\frac{1}{3}}\bar{\mathbb{A}}_{n+\frac{1}{2}}},$$

with

$$\begin{aligned}
\frac{1}{\mu}\operatorname{tr}(\mathbb{K}) &= 3\frac{K}{\mu}\frac{\partial\psi_{\mathrm{v}}}{\partial J}J + \frac{\beta}{\mu}\frac{\partial\psi_{\mathrm{ani}}}{\partial \mathrm{A}^{\hat{\theta}}{}_{\hat{\theta}}}\mathrm{A}^{\hat{\theta}}{}_{\hat{\theta}},\\
\frac{1}{\mu}\mathbb{K}^{\mathrm{d}}_{2\mathrm{D}} &= \left(\frac{\partial\psi_{\mathrm{i}}}{\partial\bar{\mathbb{A}}}\bar{\mathbb{A}}^{\top}\right)^{\mathrm{d}}_{2\mathrm{D}} + \frac{\beta}{\mu}\left(\frac{\partial\psi_{\mathrm{ani}}}{\partial\mathbb{A}}\mathbb{A}^{\top}\right)^{\mathrm{d}}_{2\mathrm{D}}\\
&= \frac{\partial\psi_{\mathrm{i}}}{\partial\bar{\mathbb{A}}_{2\mathrm{D}}}\bar{\mathbb{A}}^{\top}_{2\mathrm{D}} - \frac{1}{3}\left(\frac{\partial\psi_{\mathrm{i}}}{\partial\bar{\mathbb{A}}_{2\mathrm{D}}}:\bar{\mathbb{A}}_{2\mathrm{D}} + \frac{\partial\psi_{\mathrm{i}}}{\partial\bar{\mathrm{A}}^{\hat{\theta}}{}_{\hat{\theta}}}\bar{\mathrm{A}}^{\hat{\theta}}{}_{\hat{\theta}} + \frac{\beta}{\mu}\frac{\partial\psi_{\mathrm{ani}}}{\partial\mathrm{A}^{\hat{\theta}}{}_{\hat{\theta}}}\mathrm{A}^{\hat{\theta}}{}_{\hat{\theta}}\right)\mathbb{I}_{2\mathrm{D}},\\
\frac{1}{\mu}(\mathbb{K}^{\mathrm{d}})^{\hat{\theta}\hat{\theta}} &= \left(\frac{\partial\psi_{\mathrm{i}}}{\partial\bar{\mathbb{A}}}\bar{\mathbb{A}}^{\top}\right)^{\mathrm{d},\hat{\theta}\hat{\theta}} + \frac{\beta}{\mu}\left(\frac{\partial\psi_{\mathrm{ani}}}{\partial\mathbb{A}}\mathbb{A}^{\top}\right)^{\mathrm{d},\hat{\theta}\hat{\theta}}\\
&= \frac{2}{3}\frac{\partial\psi_{\mathrm{i}}}{\partial\bar{\mathrm{A}}^{\hat{\theta}}{}_{\hat{\theta}}}\bar{\mathrm{A}}^{\hat{\theta}}{}_{\hat{\theta}} - \frac{1}{3}\frac{\partial\psi_{\mathrm{i}}}{\partial\bar{\mathbb{A}}_{2\mathrm{D}}}:\bar{\mathbb{A}}_{2\mathrm{D}} + \frac{2}{3}\frac{\beta}{\mu}\frac{\partial\psi_{\mathrm{ani}}}{\partial\mathrm{A}^{\hat{\theta}}{}_{\hat{\theta}}}\mathrm{A}^{\hat{\theta}}{}_{\hat{\theta}},
\end{aligned}$$

and

$$\begin{aligned}
\frac{\partial\psi_{\mathrm{v}}}{\partial J} &= \frac{1}{10}\left(J^4 - J^{-6}\right),\\
\frac{\partial\psi_{\mathrm{i}}}{\partial\bar{\mathbb{A}}_{2\mathrm{D}}} &= \frac{1}{9}\left(\bar{\mathbb{A}}_{2\mathrm{D}}:\bar{\mathbb{A}}_{2\mathrm{D}} + (\bar{\mathrm{A}}^{\hat{\theta}}{}_{\hat{\theta}})^2\right)^2\bar{\mathbb{A}}_{2\mathrm{D}},\\
\frac{\partial\psi_{\mathrm{i}}}{\partial\bar{\mathrm{A}}^{\hat{\theta}}{}_{\hat{\theta}}} &= \frac{1}{9}\left(\bar{\mathbb{A}}_{2\mathrm{D}}:\bar{\mathbb{A}}_{2\mathrm{D}} + (\bar{\mathrm{A}}^{\hat{\theta}}{}_{\hat{\theta}})^2\right)^2\bar{\mathrm{A}}^{\hat{\theta}}{}_{\hat{\theta}},\\
\frac{\partial\psi_{\mathrm{ani}}}{\partial\mathrm{A}^{\hat{\theta}}{}_{\hat{\theta}}} &= 2\left(\mathrm{A}^{\hat{\theta}}{}_{\hat{\theta}} - (\mathrm{A}^{\hat{\theta}}{}_{\hat{\theta}})^{-1}\right).
\end{aligned}$$

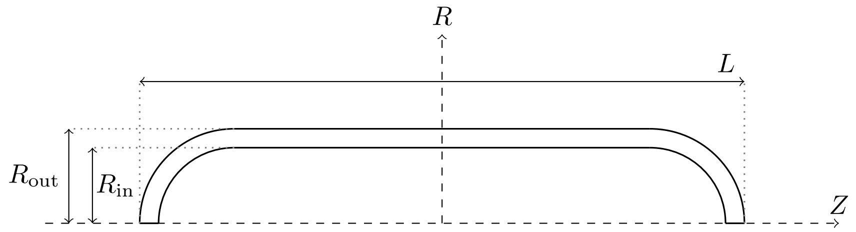

Figure 2: Half-stadium.

Since the circumferential component of the Piola–Kirchhoff stress $\mathrm{P}^{\hat{\theta}\hat{\Theta}}$ is given by a known, explicit relation (51), we have effectively reduced the problem to a two-dimensional setting by considering the "half-stadium" $B$ as the computational domain and equation (52) as the weak formulation of our problem. Thanks to the symmetry across the line $Z = 0$ we can even reduce the problem to the quarter stadium, where on the intersection of $Z = 0$ with $\partial B$ we prescribe the boundary condition $\mathrm{U}^{\hat{z}}(R, Z) = 0$ and on the intersection $R = 0$ with $\partial B$ we prescribe $\mathrm{U}^{\hat{r}}(R, Z) = 0$.

##### 3.1.2 Growth and remodeling

Using the decomposition of $\mathbb{M}$ and $\mathbb{L}_\kappa$ we rewrite the growth law (38)

$$
\begin{aligned}
&\int_B R \left( \frac{1}{\mu} \frac{\partial \mathrm{R}}{\partial (\mathbb{L}_\kappa)_{2\mathrm{D}}} \left( (\mathbb{L}_\kappa)_{n+\frac{1}{2}} \right) - \omega_{n+\frac{1}{2}} \zeta(a_n) \mathbb{I}_{2\mathrm{D}} \right) : \mathbb{V}_{2\mathrm{D}} (\mathrm{G}^{\hat{\theta}}{}_{\hat{\Theta}})_n (\det \mathbb{G}_{2\mathrm{D}})_n \, \mathrm{d}R \, \mathrm{d}Z \\
&+ \int_B R \left( \frac{1}{\mu} \frac{\partial \mathrm{R}}{\partial \mathrm{L}_\kappa^{\hat{\theta}\hat{\theta}}} \left( (\mathbb{L}_\kappa)_{n+\frac{1}{2}} \right) - \omega_{n+\frac{1}{2}} \zeta(a_n) \right) \mathrm{V}^{\hat{\theta}\hat{\theta}} (\mathrm{G}^{\hat{\theta}}{}_{\hat{\Theta}})_n (\det \mathbb{G}_{2\mathrm{D}})_n \, \mathrm{d}R \, \mathrm{d}Z \\
&= \int_B R \zeta(a_n) \frac{1}{\mu} (\mathbb{M}_{2\mathrm{D}} \mathbb{G}_{2\mathrm{D}}^{-\top})_{n+\frac{1}{2}} (\mathbb{G}_{2\mathrm{D}}^{\top})_n : \mathbb{V}_{2\mathrm{D}} (\mathrm{G}^{\hat{\theta}}{}_{\hat{\Theta}})_{n+\frac{1}{2}} (\det \mathbb{G}_{2\mathrm{D}})_{n+\frac{1}{2}} \, \mathrm{d}R \, \mathrm{d}Z
\end{aligned} \tag{56}
$$

$$
+ \int_B R \zeta(a_n) \frac{1}{\mu} (\mathrm{M}^{\hat{\theta}\hat{\theta}} (\mathrm{G}^{\hat{\theta}}{}_{\hat{\Theta}})^{-1})_{n+\frac{1}{2}} (\mathrm{G}^{\hat{\theta}}{}_{\hat{\Theta}})_n \mathrm{V}^{\hat{\theta}\hat{\theta}} (\mathrm{G}^{\hat{\theta}}{}_{\hat{\Theta}})_{n+\frac{1}{2}} (\det \mathbb{G}_{2\mathrm{D}})_{n+\frac{1}{2}} \, \mathrm{d}R \, \mathrm{d}Z, \tag{57}
$$

$$
\int_B R \operatorname{tr} \left( (\mathbb{L}_\kappa)_{n+\frac{1}{2}} \right) (\mathrm{G}^{\hat{\theta}}{}_{\hat{\Theta}})_n (\det \mathbb{G}_{2\mathrm{D}})_n \, \mathrm{d}R \, \mathrm{d}Z = \int_B R \frac{\Omega}{2\pi |B|_R} \, \mathrm{d}R \, \mathrm{d}Z, \tag{58}
$$

where one factor $2\pi$ comes from the cylindrical integration and the $R$-weighted area of the stadium is given by

$$
|B|_R := \int_B R \, \mathrm{d}R \, \mathrm{d}Z.
$$

Thanks to the symmetry across the plane $Z = 0$ we integrate in the code over the quarter-stadium and hence divide by $4\pi$ instead of $2\pi$. The tensors are given by

$$
\begin{aligned}
(\mathbb{L}_\kappa)_{n+\frac{1}{2}} &:= \frac{\mathbb{G}_{n+\frac{1}{2}} - \mathbb{G}_n}{\tau} \mathbb{G}_n^{-1}, \\
\frac{1}{\mu} (\mathbb{M}_{2\mathrm{D}})_{n+\frac{1}{2}} &:= -p_{n+\frac{1}{2}} \iota_{n+\frac{1}{2}} \mathbb{I}_{2\mathrm{D}} + \left[ \frac{1}{\mu} \mathbb{M}_{2\mathrm{D}}^{\mathrm{d}} \right]_{\mathbb{A} = (\iota_{n+\frac{1}{2}})^{\frac{1}{3}} \bar{\mathbb{A}}_{n+\frac{1}{2}}}, \\
\frac{1}{\mu} (\mathrm{M}^{\hat{\theta}\hat{\theta}})_{n+\frac{1}{2}} &:= -p_{n+\frac{1}{2}} \iota_{n+\frac{1}{2}} + \left[ \frac{1}{\mu} (\mathrm{M}^{\mathrm{d}})^{\hat{\theta}\hat{\theta}} \right]_{\mathbb{A} = (\iota_{n+\frac{1}{2}})^{\frac{1}{3}} \bar{\mathbb{A}}_{n+\frac{1}{2}}},
\end{aligned}
$$

where

$$
\begin{aligned}
\frac{1}{\mu}\frac{\partial \mathrm{R}}{\partial(\mathbb{L}_\kappa)_{2\mathrm{D}}} &= \frac{\kappa}{\mu}\frac{\partial \mathrm{R}_\mathrm{v}}{\partial \mathrm{l}_\kappa}\mathbb{I}_{2\mathrm{D}} + \frac{\eta}{\mu}\frac{\partial \mathrm{R}_\mathrm{i}}{\partial(\mathbb{L}^\mathrm{d}_\kappa)_{2\mathrm{D}}},\\
\frac{1}{\mu}\frac{\partial \mathrm{R}}{\partial \mathrm{L}_\kappa^{\hat\theta\hat\theta}} &= \frac{\kappa}{\mu}\frac{\partial \mathrm{R}_\mathrm{v}}{\partial \mathrm{l}_\kappa} + \frac{\eta}{\mu}\frac{\partial \mathrm{R}_\mathrm{i}}{\partial(\mathrm{L}^\mathrm{d}_\kappa)^{\hat\theta\hat\theta}},\\
\frac{1}{\mu}\mathbb{M}^\mathrm{d}_{2\mathrm{D}} &= \left(\bar{\mathbb{A}}^\top\frac{\partial\psi_\mathrm{i}}{\partial\bar{\mathbb{A}}}\right)^\mathrm{d}_{2\mathrm{D}} + \frac{\beta}{\mu}\left(\mathbb{A}^\top\frac{\partial\psi_\mathrm{ani}}{\partial\mathbb{A}}\right)^\mathrm{d}_{2\mathrm{D}}\\
&= \bar{\mathbb{A}}^\top_{2\mathrm{D}}\frac{\partial\psi_\mathrm{i}}{\partial\bar{\mathbb{A}}_{2\mathrm{D}}} - \frac{1}{3}\left(\frac{\partial\psi_\mathrm{i}}{\partial\bar{\mathbb{A}}_{2\mathrm{D}}} : \bar{\mathbb{A}}_{2\mathrm{D}} + \frac{\partial\psi_\mathrm{i}}{\partial\bar{\mathrm{A}}^{\hat\theta}{}_{\hat\theta}}\bar{\mathrm{A}}^{\hat\theta}{}_{\hat\theta} + \frac{\beta}{\mu}\frac{\partial\psi_\mathrm{ani}}{\partial\mathrm{A}^{\hat\theta}{}_{\hat\theta}}\mathrm{A}^{\hat\theta}{}_{\hat\theta}\right)\mathbb{I}_{2\mathrm{D}},\\
\frac{1}{\mu}(\mathbb{M}^\mathrm{d})^{\hat\theta\hat\theta} &= \left(\bar{\mathbb{A}}^\top\frac{\partial\psi_\mathrm{i}}{\partial\bar{\mathbb{A}}}\right)^{\mathrm{d},\hat\theta\hat\theta} + \frac{\beta}{\mu}\left(\mathbb{A}^\top\frac{\partial\psi_\mathrm{ani}}{\partial\mathbb{A}}\right)^{\mathrm{d},\hat\theta\hat\theta}\\
&= \frac{2}{3}\frac{\partial\psi_\mathrm{i}}{\partial\bar{\mathrm{A}}^{\hat\theta}{}_{\hat\theta}}\bar{\mathrm{A}}^{\hat\theta}{}_{\hat\theta} - \frac{1}{3}\frac{\partial\psi_\mathrm{i}}{\partial\bar{\mathbb{A}}_{2\mathrm{D}}} : \bar{\mathbb{A}}_{2\mathrm{D}} + \frac{2}{3}\frac{\beta}{\mu}\frac{\partial\psi_\mathrm{ani}}{\partial\mathrm{A}^{\hat\theta}{}_{\hat\theta}}\mathrm{A}^{\hat\theta}{}_{\hat\theta},
\end{aligned}
$$

and

$$
\begin{aligned}
\frac{\partial \mathrm{R}_\mathrm{v}}{\partial \mathrm{l}_\kappa} &= \mathrm{tr}(\mathbb{L}_\kappa) = \mathrm{tr}(\mathbb{L}_\kappa)_{2\mathrm{D}} + \mathrm{L}_\kappa^{\hat\theta\hat\theta},\\
\frac{\partial \mathrm{R}_\mathrm{i}}{\partial(\mathbb{L}^\mathrm{d}_\kappa)_{2\mathrm{D}}} &= 2(\mathbb{L}^\mathrm{d}_\kappa)_{2\mathrm{D}} = 2(\mathbb{L}_\kappa)_{2\mathrm{D}} - 2\frac{\mathrm{tr}(\mathbb{L}_\kappa)_{2\mathrm{D}} + \mathrm{L}_\kappa^{\hat\theta\hat\theta}}{3}\mathbb{I}_{2\mathrm{D}},\\
\frac{\partial \mathrm{R}_\mathrm{i}}{\partial(\mathrm{L}^\mathrm{d}_\kappa)^{\hat\theta\hat\theta}} &= 2(\mathrm{L}^\mathrm{d}_\kappa)^{\hat\theta\hat\theta} = \frac{4}{3}\mathrm{L}_\kappa^{\hat\theta\hat\theta} - 2\frac{\mathrm{tr}(\mathbb{L}_\kappa)_{2\mathrm{D}}}{3},\\
\frac{\partial\psi_\mathrm{i}}{\partial\bar{\mathbb{A}}_{2\mathrm{D}}} &= \frac{1}{9}\left(\bar{\mathbb{A}}_{2\mathrm{D}} : \bar{\mathbb{A}}_{2\mathrm{D}} + (\bar{\mathrm{A}}^{\hat\theta}{}_{\hat\theta})^2\right)^2\bar{\mathbb{A}}_{2\mathrm{D}},\\
\frac{\partial\psi_\mathrm{i}}{\partial\bar{\mathrm{A}}^{\hat\theta}{}_{\hat\theta}} &= \frac{1}{9}\left(\bar{\mathbb{A}}_{2\mathrm{D}} : \bar{\mathbb{A}}_{2\mathrm{D}} + (\bar{\mathrm{A}}^{\hat\theta}{}_{\hat\theta})^2\right)^2\bar{\mathrm{A}}^{\hat\theta}{}_{\hat\theta},\\
\frac{\partial\psi_\mathrm{ani}}{\partial\mathrm{A}^{\hat\theta}{}_{\hat\theta}} &= 2\left(\mathrm{A}^{\hat\theta}{}_{\hat\theta} - (\mathrm{A}^{\hat\theta}{}_{\hat\theta})^{-1}\right).
\end{aligned}
$$

###### 3.1.3 Eikonal equation

The sought function $a = a(R, Z)$ satisfies

$$
\varepsilon\int_B(\mathbb{C}^{-1}_{2\mathrm{D}})_{n+\frac{1}{2}}\nabla a_{n+1} : \nabla\varphi\,\mathrm{d}R\,\mathrm{d}Z + \int_B|(\mathbb{F}^{-\top}_{2\mathrm{D}})_{n+\frac{1}{2}}\nabla a_{n+1}|\varphi\,\mathrm{d}R\,\mathrm{d}Z = \int_B\varphi\,\mathrm{d}R\,\mathrm{d}Z,
$$

$$
a_{n+1} = 0, \quad \text{on the midplane of } \mathcal{B}.
$$

##### 3.1.4 ALE displacement

The ALE displacement $\vec{\phi}$ is subject to the same constraint as the physical displacement $\vec{U}$, i.e.

$$\vec{\phi} = \begin{bmatrix} \phi^{\hat{r}}(R,Z) \\ 0 \\ \phi^{\hat{z}}(R,Z) \end{bmatrix},$$

and its differential is then given by

$$\nabla\vec{\phi} = \begin{bmatrix} \frac{\partial \phi^{\hat{r}}}{\partial R} & 0 & \frac{\partial \phi^{\hat{r}}}{\partial Z} \\ 0 & \frac{\phi^{\hat{r}}}{R} & 0 \\ \frac{\partial \phi^{\hat{z}}}{\partial R} & 0 & \frac{\partial \phi^{\hat{z}}}{\partial Z} \end{bmatrix}. \tag{59}$$

Using the structure of the tensors the $\vec{\phi}$ is solution to the following problem

$$\vec{\phi} \in \operatorname{ArgMin} \frac{1}{2} \int_B \left( R|(\mathbb{I}_{2D} + \nabla\vec{\phi}_{2D})\mathbb{G}_{2D}^{-1} - \mathbb{I}_{2D}|^2 + \left((1+\phi^{\hat{r}})(\mathrm{G}^{\hat{\theta}}{}_{\hat{\Theta}})^{-1} - 1\right)^2 \right) \mathrm{G}^{\hat{\theta}}{}_{\hat{\Theta}} (\det \mathbb{G}_{2D})\, \mathrm{d}R\, \mathrm{d}Z.$$

### 4 Appendix

#### 4.1 A note on unit conversions

Since we use $\mu$m as a length unit in the implementation, we need to convert the other variables appropriately. For example, the unit MPa is converted as follows

$$\mathrm{MPa} = 10^6\,\mathrm{kg}\cdot\mathrm{m}^{-1}\cdot\mathrm{s}^{-2} = \mathrm{kg}\cdot\mu\mathrm{m}^{-1}\cdot\mathrm{s}^{-2}. \tag{60}$$

$$\mathrm{MPa}\cdot\mu\mathrm{m}^2 = 10^{-12}\,\mathrm{MPa}\cdot 10^{12}\,\mu\mathrm{m}^2 = 10^{-6}\,\mathrm{Pa}\cdot\mathrm{m}^2 = \mu\mathrm{N}. \tag{61}$$

$$\mu\mathrm{N}\cdot\mu\mathrm{m}^{-1} = 10^3\,\mathrm{pN}\cdot\mathrm{nm}^{-1} \tag{62}$$

At 298 K we have

$$10\,\mathrm{MPa} \approx 4\,\mathrm{Mol}\cdot\mathrm{kg}^{-1}. \tag{63}$$

#### 4.2 Elastic moduli of quadratic envelope

For linear, elastic, and transverse isotropic material with structural tensor $\vec{m}_{\mathrm{R}} \otimes \vec{m}_{\mathrm{R}} = \vec{e}_{\hat{\theta}} \otimes \vec{e}_{\hat{\theta}}$ we have in the Voigt notatino

$$\begin{bmatrix} \mathrm{T}^{\hat{r}\hat{r}} \\ \mathrm{T}^{\hat{z}\hat{z}} \\ \mathrm{T}^{\hat{\theta}\hat{\theta}} \\ \mathrm{T}^{\hat{z}\hat{\theta}} \\ \mathrm{T}^{\hat{r}\hat{\theta}} \\ \mathrm{T}^{\hat{r}\hat{z}} \end{bmatrix} = \begin{bmatrix} \mathrm{C}_{11} & \mathrm{C}_{12} & \mathrm{C}_{13} & 0 & 0 & 0 \\ \mathrm{C}_{12} & \mathrm{C}_{11} & \mathrm{C}_{13} & 0 & 0 & 0 \\ \mathrm{C}_{13} & \mathrm{C}_{13} & \mathrm{C}_{33} & 0 & 0 & 0 \\ 0 & 0 & 0 & \mathrm{C}_{44} & 0 & 0 \\ 0 & 0 & 0 & 0 & \mathrm{C}_{44} & 0 \\ 0 & 0 & 0 & 0 & 0 & \frac{1}{2}(\mathrm{C}_{11} - \mathrm{C}_{12}) \end{bmatrix} \begin{bmatrix} \varepsilon^{\hat{r}\hat{r}} \\ \varepsilon^{\hat{z}\hat{z}} \\ \varepsilon^{\hat{\theta}\hat{\theta}} \\ 2\varepsilon^{\hat{z}\hat{\theta}} \\ 2\varepsilon^{\hat{r}\hat{\theta}} \\ 2\varepsilon^{\hat{r}\hat{z}} \end{bmatrix},$$

where the tensor of elastic constant has the form

$$\begin{bmatrix} \mathrm{c}_1 & \mathrm{c}_1+\mathrm{c}_2 & \mathrm{c}_1+\mathrm{c}_2+\mathrm{c}_4 & 0 & 0 & 0 \\ \mathrm{c}_1+\mathrm{c}_2 & \mathrm{c}_1 & \mathrm{c}_1+\mathrm{c}_2+\mathrm{c}_4 & 0 & 0 & 0 \\ \mathrm{c}_1+\mathrm{c}_2+\mathrm{c}_4 & \mathrm{c}_1+\mathrm{c}_2+\mathrm{c}_4 & \mathrm{c}_1+\mathrm{c}_3+2(\mathrm{c}_4+\mathrm{c}_5) & 0 & 0 & 0 \\ 0 & 0 & 0 & \dfrac{\mathrm{c}_5-\mathrm{c}_2}{2} & 0 & 0 \\ 0 & 0 & 0 & 0 & \dfrac{\mathrm{c}_5-\mathrm{c}_2}{2} & 0 \\ 0 & 0 & 0 & 0 & 0 & \dfrac{-\mathrm{c}_2}{2} \end{bmatrix}$$

and the constants $c_i$, $i = 1,\ldots,5$, are given in terms of the Helmholtz free energy $\psi$ as

$$\begin{aligned} \mathrm{c}_1 &= 4\left(\frac{\partial^2\psi}{\partial I_1\partial I_1} + 4\frac{\partial^2\psi}{\partial I_2\partial I_2} + \frac{\partial^2\psi}{\partial I_3\partial I_3} + 4\frac{\partial^2\psi}{\partial I_2\partial I_3} + 2\frac{\partial^2\psi}{\partial I_1\partial I_3} + 4\frac{\partial^2\psi}{\partial I_1\partial I_2}\right), \\ \mathrm{c}_2 &= 4\left(\frac{\partial\psi}{\partial I_2} + \frac{\partial\psi}{\partial I_3}\right), \\ \mathrm{c}_3 &= 4\left(\frac{\partial^2\psi}{\partial J_4\partial J_4} + 4\frac{\partial^2\psi}{\partial J_5\partial J_5} + 4\frac{\partial^2\psi}{\partial J_4\partial J_5}\right), \\ \mathrm{c}_4 &= 4\left(\frac{\partial^2\psi}{\partial I_1\partial J_4} + 2\frac{\partial^2\psi}{\partial I_1\partial J_5} + 2\frac{\partial^2\psi}{\partial I_2\partial J_4} + 4\frac{\partial^2\psi}{\partial I_2\partial J_5} + \frac{\partial^2\psi}{\partial I_3\partial J_4} + 2\frac{\partial^2\psi}{\partial I_3\partial J_5}\right), \\ \mathrm{c}_5 &= 4\frac{\partial\psi}{\partial J_5} \end{aligned}$$

in terms of the invariants $I_j$, $j = 1,\ldots,5$

$$I_1 = \operatorname{tr}\mathbb{C}_{\mathrm{el}}, \quad I_2 = \operatorname{tr}(\operatorname{Cof}\mathbb{C}_{\mathrm{el}}), \quad I_3 = \det\mathbb{C}_{\mathrm{el}}, \quad J_4 = \mathbb{C}_{\mathrm{el}} : \vec{m}_{\mathrm{R}}\otimes\vec{m}_{\mathrm{R}}, \quad J_5 = \mathbb{C}_{\mathrm{el}}^2 : \vec{m}_{\mathrm{R}}\otimes\vec{m}_{\mathrm{R}}.$$

The terms appearing in the free energy $\psi$ can be expressed as

$$J = I_3^{\frac{1}{2}}, \quad |\bar{\mathbb{A}}|^2 = \operatorname{tr}\bar{\mathbb{C}}_{\mathrm{el}} = I_3^{-\frac{1}{3}} I_1, \quad |\mathbb{A}\vec{m}_{\mathrm{R}}|^2 = \mathbb{C}_{\mathrm{el}} : \vec{m}_{\mathrm{R}}\otimes\vec{m}_{\mathrm{R}} = J_4.$$

The energy contributions are then

$$\psi_{\mathrm{v}}(I_3) = \frac{1}{50}\big(I_3^{\frac{5}{2}} + I_3^{-\frac{5}{2}} - 2\big), \quad \psi_{\mathrm{i}}(I_1, I_3) = \frac{1}{54}\left(I_3^{-1}I_1^3 - 27\right), \quad \psi_{\mathrm{ani}}(J_4) = J_4 - 1 - \ln J_4,$$

and hence evaluation at $\mathbb{C}_{\mathrm{el}} = \mathbb{I}$ yields

$$\begin{aligned} &\frac{\partial\psi_{\mathrm{v}}}{\partial I_3} = \frac{1}{20}\left(I_3^{\frac{3}{2}} - I_3^{-\frac{7}{2}}\right) = 0, \quad \frac{\partial^2\psi_{\mathrm{v}}}{\partial I_3\partial I_3} = \frac{1}{20}\left(\frac{3}{2}I_3^{\frac{1}{2}} - \frac{7}{2}I_3^{-\frac{9}{2}}\right) = \frac{1}{4}, \\ &\frac{\partial\psi_{\mathrm{i}}}{\partial I_1} = \frac{1}{18}I_1^2 I_3^{-1} = \frac{1}{2} \quad \frac{\partial\psi_{\mathrm{i}}}{\partial I_3} = -\frac{1}{54}I_1^3 I_3^{-2} = -\frac{1}{2}, \\ &\frac{\partial^2\psi_{\mathrm{i}}}{\partial I_1\partial I_1} = \frac{1}{9}I_1 I_3^{-1} = \frac{1}{3}, \quad \frac{\partial^2\psi_{\mathrm{i}}}{\partial I_1\partial I_3} = -\frac{1}{18}I_1^2 I_3^{-2} = -\frac{1}{2}, \quad \frac{\partial^2\psi_{\mathrm{i}}}{\partial I_3\partial I_3} = \frac{1}{27}I_1^3 I_3^{-3} = 1, \\ &\frac{\partial\psi_{\mathrm{ani}}}{\partial J_4} = 1 - J_4^{-1} = 0, \quad \frac{\partial^2\psi_{\mathrm{ani}}}{\partial J_4\partial J_4} = J_4^{-2} = 1. \end{aligned}$$

In terms of the elastic moduli we have

$$
\begin{aligned}
\mathrm{c}_1 &= 4\left(\frac{\mu}{3} + \frac{K}{4} + \mu - \mu\right) = K + \frac{4}{3}\mu, \\
\mathrm{c}_2 &= -2\mu, \\
\mathrm{c}_3 &= 4\beta, \\
\mathrm{c}_4 &= 0, \\
\mathrm{c}_5 &= 0,
\end{aligned}
$$

and hence the tensor of elastic constants reads

$$
\mathcal{C} = \begin{bmatrix}
K + \frac{4}{3}\mu & K - \frac{2}{3}\mu & K - \frac{2}{3}\mu & 0 & 0 & 0 \\
 & K + \frac{4}{3}\mu & K - \frac{2}{3}\mu & 0 & 0 & 0 \\
 & & K + \frac{4}{3}\mu + 4\beta & 0 & 0 & 0 \\
 & & & \mu & 0 & 0 \\
 & & & & \mu & 0 \\
 & & & & & \mu
\end{bmatrix}.
$$

For computing the Young moduli and Poisson ratio we invert the tensor to obtain the so-called compliance tensor. We have

$$
\begin{aligned}
\Delta &= (\mathrm{C}_{11} - \mathrm{C}_{12})[(\mathrm{C}_{11} + \mathrm{C}_{12})\mathrm{C}_{33} - 2\mathrm{C}_{13}\mathrm{C}_{13}] \\
&= 2\mu\left(\left(2K + \frac{2}{3}\mu\right)\left(K + \frac{4}{3}\mu + 4\beta\right) - 2\left(K - \frac{2}{3}\mu\right)^2\right) \\
&= 2\mu\left(2K^2 + \frac{8}{9}\mu^2 + \frac{10}{3}K\mu + 8K\beta + \frac{8}{3}\mu\beta - 2K^2 - \frac{8}{9}\mu^2 + \frac{8}{3}K\mu\right) \\
&= \frac{4}{3}\mu\left(9K\mu + 12K\beta + 4\mu\beta\right),
\end{aligned}
$$

and

$$
\mathcal{C}^{-1} = \frac{1}{\Delta}\begin{bmatrix}
\mathrm{C}_{11}\mathrm{C}_{33} - \mathrm{C}_{13}^2 & \mathrm{C}_{13}^2 - \mathrm{C}_{12}\mathrm{C}_{33} & (\mathrm{C}_{12} - \mathrm{C}_{11})\mathrm{C}_{13} & 0 & 0 & 0 \\
 & \mathrm{C}_{11}\mathrm{C}_{33} - \mathrm{C}_{13}^2 & (\mathrm{C}_{12} - \mathrm{C}_{11})\mathrm{C}_{13} & 0 & 0 & 0 \\
 & & \mathrm{C}_{11}^2 - \mathrm{C}_{12}^2 & 0 & 0 & 0 \\
 & & & \frac{\Delta}{\mathrm{C}_{44}} & 0 & 0 \\
 & & & & \frac{\Delta}{\mathrm{C}_{44}} & 0 \\
 & & & & & \frac{2\Delta}{\mathrm{C}_{11} - \mathrm{C}_{12}}
\end{bmatrix}.
$$

Since

$$
\begin{aligned}
C_{11}C_{33} - C_{13}^2 &= \left(K + \frac{4}{3}\mu\right)\left(K + \frac{4}{3}\mu + 4\beta\right) - \left(K - \frac{2}{3}\mu\right)^2 \\
&= K^2 + \frac{16}{9}\mu^2 + \frac{8}{3}K\mu + 4K\beta + \frac{16}{3}\mu\beta - K^2 + \frac{4}{3}K\mu - \frac{4}{9}\mu^2 \\
&= \frac{4}{3}\left(\mu^2 + 3K\mu + 3K\beta + 4\mu\beta\right), \\
C_{13}^2 - C_{12}C_{33} &= \left(K - \frac{2}{3}\mu\right)^2 - \left(K - \frac{2}{3}\mu\right)\left(K + \frac{4}{3}\mu + 4\beta\right) \\
&= \left(K - \frac{2}{3}\mu\right)\left(K - \frac{2}{3}\mu - K - \frac{4}{3}\mu - 4\beta\right) \\
&= -\left(K - \frac{2}{3}\mu\right)(2\mu + 4\beta) \\
&= \frac{2}{3}\left(2\mu^2 - 3K\mu - 6K\beta + 4\mu\beta\right), \\
(C_{12} - C_{11})C_{13} &= -2\mu\left(K - \frac{2}{3}\mu\right), \\
C_{11}^2 - C_{12}^2 &= 4\mu\left(K + \frac{1}{3}\mu\right),
\end{aligned}
$$

and

$$
\mathcal{C}^{-1} = \begin{bmatrix}
\frac{1}{E_{\hat r}} & -\frac{\nu_{\hat z\hat r}}{E_{\hat r}} & -\frac{\nu_{\hat\theta\hat r}}{E_{\hat\theta}} & 0 & 0 & 0 \\
 & \frac{1}{E_{\hat r}} & -\frac{\nu_{\hat\theta\hat r}}{E_{\hat\theta}} & 0 & 0 & 0 \\
 & & \frac{1}{E_{\hat\theta}} & 0 & 0 & 0 \\
 & & & \frac{1}{\mu} & 0 & 0 \\
 & & & & \frac{1}{\mu} & 0 \\
 & & & & & \frac{2(1+\nu_{\hat r\hat z})}{E_{\hat r}}
\end{bmatrix},
$$

the sought relations read

$$
\begin{aligned}
E_{\hat r} = E_{\hat z} &= \frac{\Delta}{C_{11}C_{33} - C_{13}^2} = \mu\frac{9K\mu + 12K\beta + 4\mu\beta}{\mu^2 + 3K\mu + 3K\beta + 4\mu\beta} = \mu\frac{9K\mu + 4\beta(3K+\mu)}{\mu(3K+\mu) + \beta(3K+4\mu)}, \\
E_{\hat\theta} &= \frac{\Delta}{C_{11}^2 - C_{12}^2} = \frac{9K\mu + 12K\beta + 4\mu\beta}{3K+\mu} = \frac{9K\mu + 4\beta(3K+\mu)}{3K+\mu} = \frac{9K\mu}{3K+\mu} + 4\beta, \\
\nu_{\hat\theta\hat z} &= \frac{1}{2}\frac{3K - 2\mu}{3K+\mu}.
\end{aligned}
$$

#### 4.3 Computing volume and area

We can compute volume enclosed by the cell wall via Stokes formula. In the last step the factor $2 \cdot 2\pi$ pops up, where $2\pi$ comes from integration over rotated surface and2 from the symmetry across the line $Z = 0$. Here $\Gamma_{\text{in}}$ denotes $(\partial B)_{\text{in}}$ intersected with $Z > 0$. Hence we have

$$\begin{aligned}\text{Volume} &= \int_{\text{deff.cell}} 1 \,\mathrm{d}v = \int_{\text{deff.cell}} \operatorname{div} \vec{f} \,\mathrm{d}v = \int_{(\partial \mathcal{B}_t)_{\text{in}}} \vec{f} \cdot \vec{n} \,\mathrm{d}s \\ &= \int_{(\partial \mathcal{B})_{\text{in}}} \vec{f} \cdot (\operatorname{Cof} \mathbb{F}) \vec{n}_{\text{R}} \,\mathrm{d}S = 4\pi \int_{\Gamma_{\text{in}}} R\vec{f} \cdot (\operatorname{Cof} \mathbb{F}) \vec{n}_{\text{R}} \,\mathrm{d}S,\end{aligned}$$

where

$$\vec{f} = \begin{bmatrix} 0 \\ 0 \\ z \end{bmatrix} = \begin{bmatrix} 0 \\ 0 \\ z(Z,R) \end{bmatrix} = \begin{bmatrix} 0 \\ 0 \\ Z + \mathrm{U}^{\hat{z}}(R,Z) \end{bmatrix}.$$

Similarly for the area enclosed by $(\partial B_t)_{\text{in}}$ we have

$$\begin{aligned}\text{Area} &= \int_{\text{deff.plane}} 1 \,\mathrm{d}R \,\mathrm{d}Z = \int_{\text{deff.plane}} \operatorname{div} \vec{g} \,\mathrm{d}R \,\mathrm{d}Z = \int_{(\partial B_t)_{\text{in}}} \vec{g} \cdot \vec{n} \,\mathrm{d}l \\ &= \int_{(\partial B)_{\text{in}}} \vec{g} \cdot (\operatorname{Cof} \mathbb{F}_{2\text{D}}) \vec{n}_{\text{R}} \,\mathrm{d}L = 4 \int_{\Gamma_{\text{in}}} \vec{g} \cdot (\operatorname{Cof} \mathbb{F}_{2\text{D}}) \vec{n}_{\text{R}} \,\mathrm{d}L,\end{aligned}$$

where

$$\vec{g} = \begin{bmatrix} 0 \\ z \end{bmatrix} = \begin{bmatrix} 0 \\ z(Z,R) \end{bmatrix} = \begin{bmatrix} 0 \\ Z + \mathrm{U}^{\hat{z}}(R,Z) \end{bmatrix}.$$

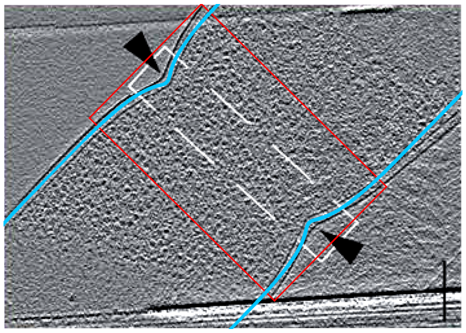

Figure 3: Data taken from Navarro et al. [2022]

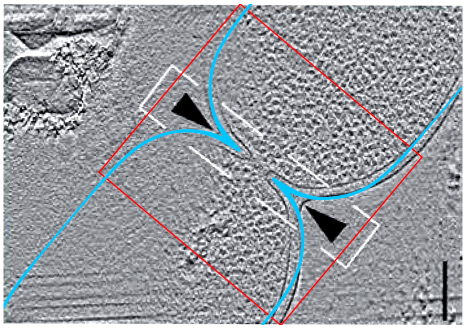

Figure 4: Data taken from Navarro et al. [2022]

##### 4.4 Processing of microscopic data

Here we show full microscopic images from which we cut the division site. The computed PG layer (in blue) is scaled to the real bacterium in each picture such that the center of the septum, its orientation and width of the bacterium agree. As on each micrograph different individua (having different widths) are shown, such a scaling is the only logical way of comparison of shapes.

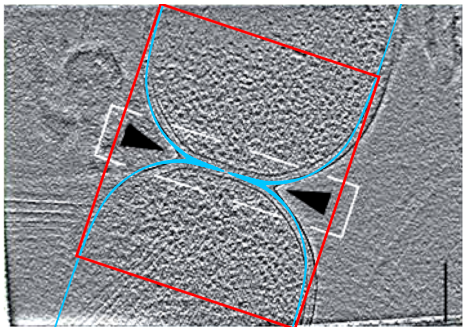

Figure 5: Data taken from Navarro et al. [2022]

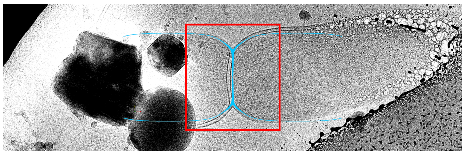

Figure 6: ΔenvC-mutant

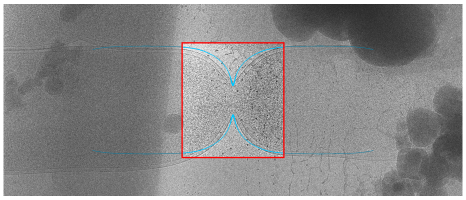

Figure 7: ftsN-ΔSPOR-mutant
